# Molecular organization of the eukaryotic secretory pathway

**DOI:** 10.64898/2026.07.28.741320

**Authors:** Courtney J. Mycroft-West, Neville B-y Yee, Miron A. Leanca, Michele C. Darrow, Liang Wu

## Abstract

The eukaryotic secretory pathway is an essential subcellular network comprised of the endoplasmic reticulum (ER) and Golgi organelles. It mediates the trafficking of protein, lipid and other biomolecular cargoes throughout the cell, and is also the site of enzymatic modifications such as glycosylation that are vital for biological function. Complex interactions between resident functional proteins and cargo biomolecules drive myriad activities within the secretory pathway, but how these components organize within native milieu is still poorly understood. Here, we used cryo-electron tomography to survey the HeLa secretory pathway, revealing detailed features of its organellar structure. By quantitatively mapping protein distributions, we find that interactions within the HeLa secretory pathway are dominated by small dimeric complexes, stochastically distributed across both ER and Golgi membranes. Our survey of secretory pathway biomolecules provides new insights into putative trafficking and modification mechanisms, and highlights links between molecular organization and function within intracellular compartments.

## Introduction

The secretory pathway is an essential multi-organellar network present in all eukaryotic cells, which is comprised of the endoplasmic reticulum (ER) and Golgi apparatus, alongside their associated transport vesicles. This network mediates the trafficking of protein, lipid, and other biomolecular cargoes within cells, moving them from initial sites of biosynthesis, to the locations ultimately required for their function. Canonically, eukaryotic secretory pathways operate in a highly directional manner, with new biomolecular cargoes first being processed in the ER, before being moved to the *cis*-Golgi *via* COPII vesicle carriers. Within the Golgi, biomolecular cargoes transit further from *cis-* to *trans-* through successive cisternae, before ultimately exiting *via* the trans-Golgi network (TGN) to their final intracellular or extracellular destinations (**Supplemental Figure 1**)^1^.

During transit, secretory pathway cargoes may be acted upon by a host of resident ER and Golgi enzymes, which mediate key modification reactions such as disulfide bond formation^2^, signal peptide cleavage^3^, and glycosylation^4^. In particular, the secretory pathway represents a principal hub for complex glycan processing in eukaryotic cells, with its resident glycosyltransferases, glycosidases and other enzymes (*e.g.* sulfotransferases, epimerases) forming a significant proportion of proteins within both ER and Golgi organelles^5,6^. Protein and lipid-linked glycans produced in the secretory pathway influence essential biological functions, including protein folding^7^, cellular recognition^8^, pathogen interactions^9^, and mitogenic signalling^10^, highlighting the key importance of understanding their construction.

The myriad trafficking and processing functions of the secretory pathway are largely regulated by the organization and distribution of its resident proteins, which creates distinct functional repertoires across each of its ER and Golgi compartments^11^. These non-uniform protein distributions particularly influence multistep reaction pathways such as glycosylation, wherein localization of early acting enzymes to the ER/*cis-*Golgi, and late acting enzymes to the *medial-*/*trans-*Golgi, allows substrate processing to operate in tandem with trafficking^12^. Physical interplay between ER/Golgi resident glycosylation enzymes is also widespread, with the formation of numerous ‘kin-interactions’^13–23^ theorized to improve multienzyme reaction efficiencies by facilitating substrate transfer. The key importance of molecular organization in shaping secretory pathway function has prompted extensive studies by both proteomic^24–26^ and fluorescence imaging^27–29^ strategies, bringing insights into broad-scale distributions and colocalizations of proteins within the ER/Golgi. However, a detailed description of how secretory pathway constituents may interplay at the molecular level is still lacking.

Electron imaging methods have been widely used to examine structure and function of the eukaryotic secretory pathway. Of particular note, 3-dimensional electron tomography (ET) studies, pioneered by Staehlin and coworkers in fixed plastic-sectioned cells, were instrumental in revealing the detailed membrane ultrastructure of ER and Golgi organelles, enabling sophisticated models of trafficking and organellar maturation to be developed^30–36^. With the resolutions now afforded by modern focused ion-beam (FIB)-milling and cryo-ET technologies, it has become possible to directly examine biomolecules *in situ*^37^. Cryo-electron tomograms of Golgi stacks, collected from vitrified sections of the unicellular algae *Chlamydomonas reinhardtii*, revealed distinctive arrayed protein assemblies within its *trans-*cisternae^38^, and molecular structures of its COPI coat machinery^39^. More recently, cryo-ET has also been used to examine the structural basis of COPII ER exit sites within human cells^40^. Whilst these studies have intensively examined key secretory pathway structures, such as vesicles involved in transport, the nature of molecular organization within core ER/Golgi compartments remains relatively unexplored.

Here, we report a workflow for fluorescence targeted cryo-ET of the HeLa secretory pathway, enabling detailed examination of its native biomolecular organization. By exploiting the nanometer resolutions possible with cryo-ET, we conducted quantitative molecular surveys of the HeLa secretory pathway, finding that proteins within its ER and Golgi compartments are densely but stochastically organized, and form interactions dominated by small dimeric complexes. Our observations provide insight into how complex secretory pathway functions such as glycosylation may be arranged, in turn highlighting links between organization and function within the intracellular environment.

## Results

### Fluorescent labelling strategies for secretory pathway CLEM

To support efficient cryo-ET imaging of the HeLa secretory pathway, we first sought to build a correlated light and electron microscopy (CLEM) workflow, which would exploit fluorescent regions of interest to guide FIB-milling and cryo-ET acquisition^41^. Due to the central position of the Golgi apparatus within the eukaryotic secretory pathway, and its close interface with both ER and transport vesicles, we reasoned that labelling this organelle alone would be sufficient for CLEM of the broader network.

Both overexpression and chemical strategies were tested for Golgi fluorescent labeling. Initially, we explored cell transfection using plasmids encoding for GFP-tagged Golgi-resident enzymes. Distinctive peri-nuclear fluorescence was observed in cells following transfection of MGAT1-GFP^42^, MGAT5-GFP^43,44^ and EXT1/2-GFP^19^, consistent with efficient translocation of these proteins to the Golgi ribbon (**Figure 1a**). Essentially identical fluorescence patterns were also observed using a split-GFP approach^45^, wherein GFP11-tagged EXT1/2 was complemented with doxycycline-induced GFP(1-10) (**Figure 1b**; **Supplementary Figure 2**). Fluorescence intensities in each case were sufficient to support FACS enrichment of labelled cells, before seeding onto EM grids. Surprisingly, not all overexpressed Golgi enzymes displayed expected localizations. Transfection of C1GALT1-GFP^46^ or NDST1-GFP^47^ led to predominantly cytosolic fluorescence, likely reflecting a need for additional partners to enable their movement to the Golgi (*e.g.* COSMC for C1GALT1^48^; EXT2 for NDST1^49^).

Next, we examined chemical labelling of cells. Compared to overexpression, treatment of eukaryotic cells with fluorescently conjugated ceramide dyes produced rapid and quantitative labeling^50^, and was also compatible with cells already seeded onto grids, removing the need for FACS enrichment (**Figure 1c**). This highly-efficient labelling was somewhat marred by increased non-specific signal at increased dye concentrations, likely due to the spreading of ceramides throughout cellular membranes. However, despite being less specific than overexpression, we considered that the substantially greater ease of chemical labeling was more suited to rapid CLEM workflows. As such, this labeling strategy was taken forward for further sample processing.

In all cases, fluorescent cells on EM grids were vitrified by plunge-freezing, and electron transparent lamellae were fabricated using plasma FIB-milling with Argon (**Figure 1**)^51^. The fluorescent signals from labelled Golgi enabled targeting of sites for lamellae fabrication and tilt-series acquisitions, supporting automated collection of datasets containing Golgi stacks, ER membranes, and other secretory pathway features. A representative set of 57 tomograms (of 519 total), collected at 42,000x (nominal 3.05 Å/pixel) and 64,000x (1.90 Å/pixel) magnifications, are summarized in **Supplemental Figure 3** and **Supplemental Table 1**.

**Figure 1.**
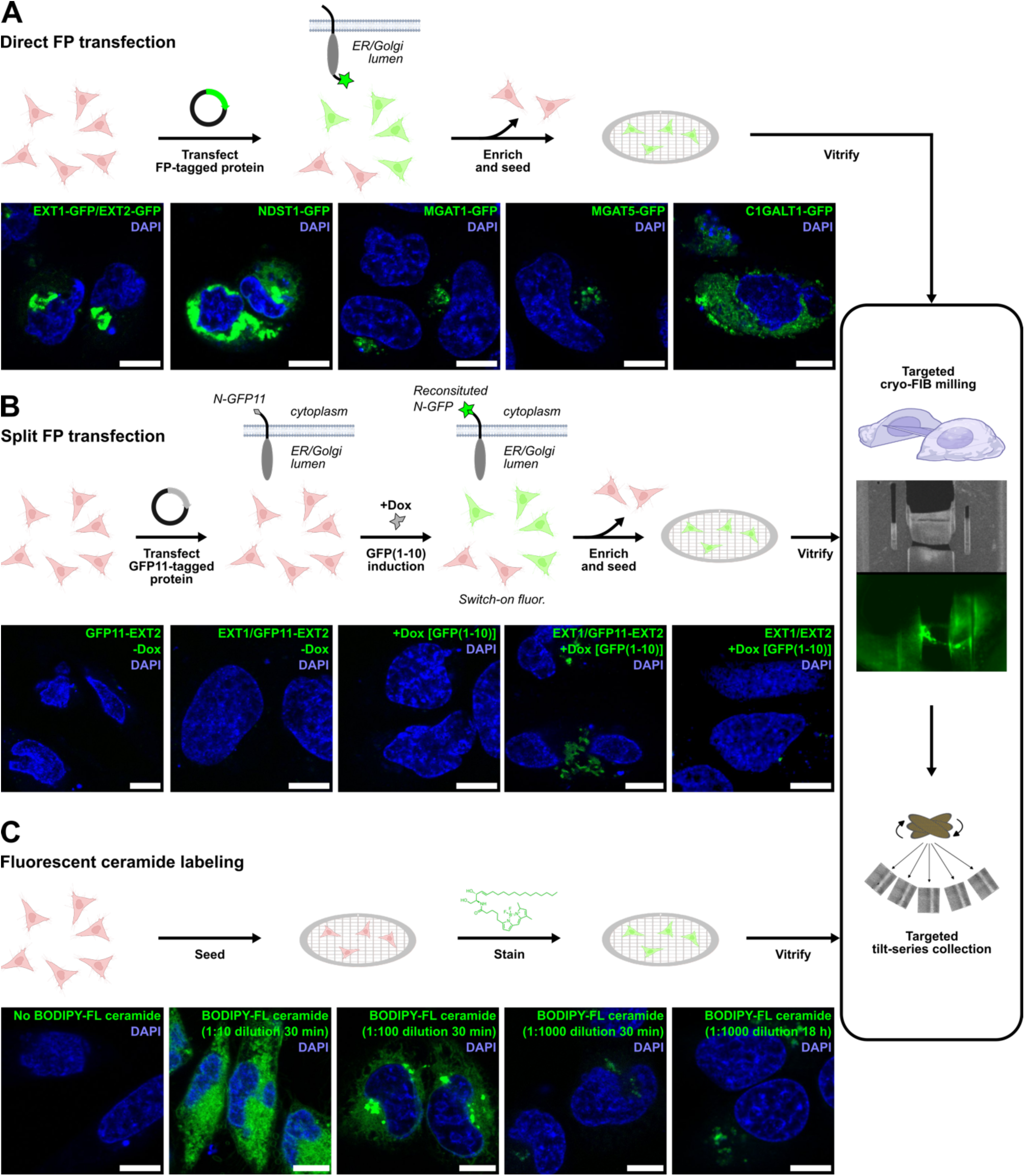
Fluorescent Golgi labeling to support CLEM of the HeLa secretory pathway. (**A**) Overexpression of GFP-tagged enzymes EXT1/2, MGAT1 and MGAT5 leads to specific perinuclear fluorescence, indicative of Golgi ribbon labelling. C1GALT1 and NDST1 overexpression produce cytoplasmic fluorescence. (**B**) Split-GFP labelling using GFP11-EXT2 shows similar labeling performance compared to direct EXT1/2-GFP overexpression. (**C**) Chemical staining using fluorescent ceramides produces rapid Golgi labelling, albeit with reduced specificity compared to overexpression. Scale bars – 10 µm.

### Ultrastructural features of the HeLa secretory pathway

Of the tomograms presented in this study (**Supplemental Figures 4**), 12 volumes were found to contain recognizable Golgi stack features (**Figure 2a**; **Supplemental Table 1**), including one volume bearing two Golgi stacks within the same field of view (Position110). Numerous other secretory pathway features were also visible, validating our strategy of Golgi labeling for CLEM of the wider network. Most volumes we reconstructed contained recognizable ER membranes and compartments (both smooth and rough ER), reflecting the widespread distribution of this organelle throughout the intracellular volume. In line with our capture of a functionally active network, COPI and COPII vesicles (including budding vesicles *e.g.* Position047, Position050) were also observed in most tomograms, which could be distinguished *via* the characteristically dense protein coats on the surface of the former^39^. Beyond core secretory pathway structures, other features of interest captured in our tomograms included: ER–mitochondria encounter structures (ERMES) postulated to be important for lipid exchange (*e.g.* Position123)^52^, numerous nuclear pore complexes near Golgi stacks, and an unusual microtubule dense array (Position006) (**Figure 2b**).

Given the distinctive morphologies that can be adopted by Golgi stacks^30–36^, and their varied structures across different cell types, we further specifically sought to examine the features of this organelle within our HeLa tomograms. To conduct ultrastructural analyses, we focused on tomograms collected at 42,000x magnification (**Supplemental Figure 3a**), which possess a larger field of view than those collected at 64,000x. Inspection of segmented membranes from Positions075, 076, 110, 113, 147, and 190 showed that HeLa Golgi stacks are typically comprised of 3-5 moderately fenestrated cisternae with highly elongated morphologies, which in some cases spanned beyond the 1.2 µm field of view of our tomograms (**Figure 2a**, **Supplemental Figure 4**). Next, we also quantitated the intra- and inter-cisternal distances adopted by Golgi stacks, moving from *cis-*Golgi cisternae (defined as side nearest to ER) to *trans*-Golgi cisternae. Pairwise nearest-distance measurements of segmented membranes in Position190 (**Figure 2c**) showed a notable bimodal distribution of intracisternal diameters within each Golgi cisterna, with peaks at ∼31-42 nm and ∼55-67 nm corresponding to thinner central regions and dilated rims, respectively (**Figure 2d**). There was a modest trend towards cisternal narrowing from *cis-* to *trans*-Golgi, with the internal diameters of central cisternal regions shifting from ∼42 nm to ∼31 nm along the Golgi stack. Similar analyses of *inter*cisternal distances showed that the separation of adjacent cisternae remained relatively consistent throughout a stack, with distances of 12-27 nm between all cisternal pairs examined (**Figure 2d**).

Taken together, our observations present a picture of highly ‘disordered’ morphologies within HeLa Golgi, which contrast markedly with the symmetrical structures that have reported in *e.g. C. reinhardtii*^38,39,53,54^. Such disordered Golgi morphologies likely reflect both the distribution of HeLa Golgi stacks into dynamic ribbon structures^55^, as well as the rapidly dividing nature of HeLa cells, wherein continuous cell cycling may preclude formation of more well-defined organelles^56^.

**Figure 2.**
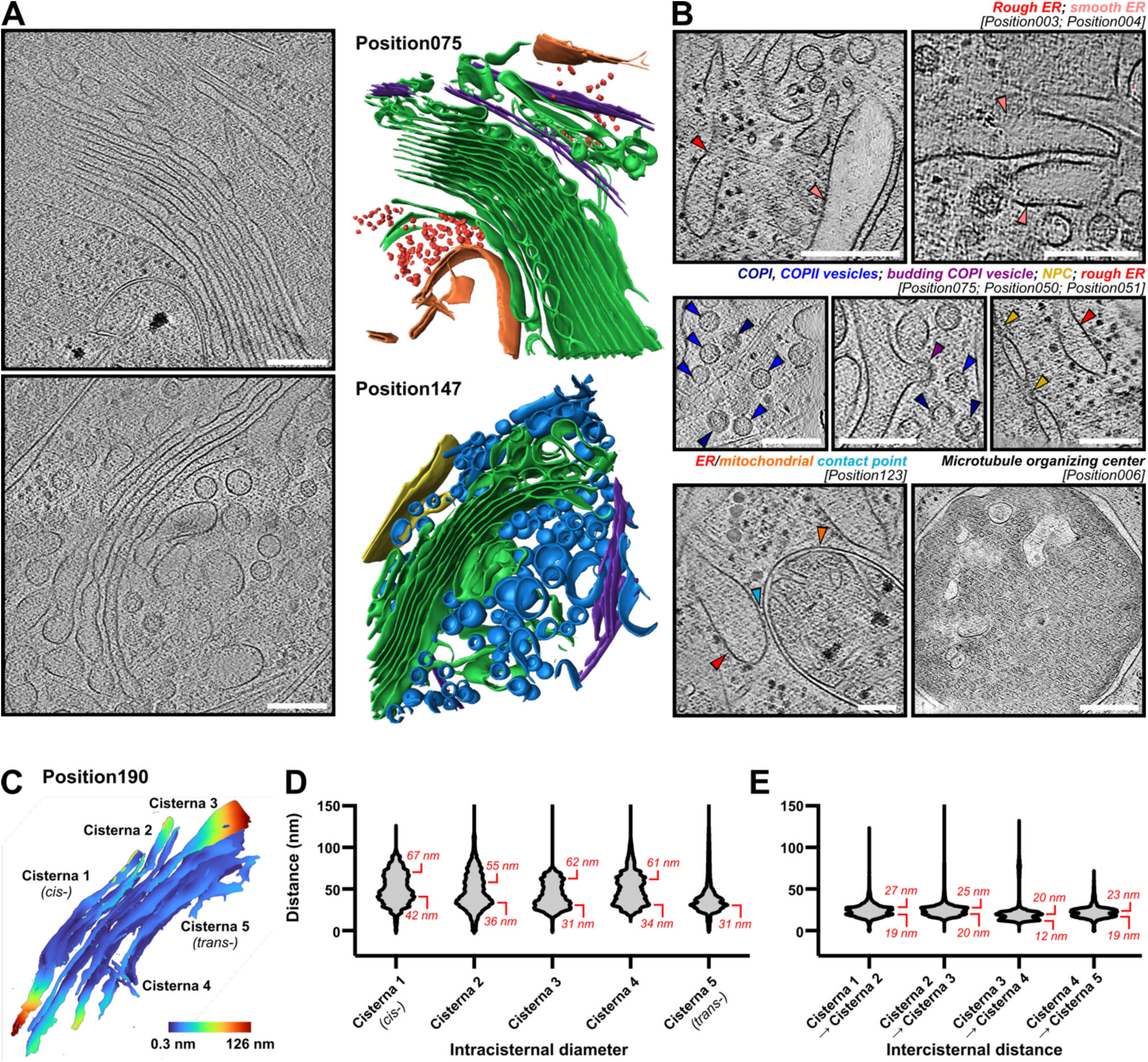
Ultrastructural features of the HeLa secretory pathway. (**A**) Tomograms of Position075 and Position113, alongside segmentation of major features. Green – Golgi; orange – mitochondrial membranes; purple – microtubules; blue – transport vesicles; yellow – nuclear membrane. (**B**) Close-up views of selected secretory pathway structures: rough and smooth ER; COPI and COPII vesicles; nuclear pore complexes; ER/mitochondrial contact sites; microtubule organizing center. (**C**) Heatmap of pairwise nearest-distance measurements between adjacent membranes in Position190. (**D**) Intracisternal widths across the cisternae of Position190, moving from cis- to trans-. (**E**) Intercisternal distances across the cisternae of Position190, moving from cis- to trans-. Scale bars – 200 nm.

### Quantitative molecular surveys of secretory pathway proteins

Close inspection of our HeLa secretory pathway tomograms revealed numerous densities across the luminal face of each ER and Golgi membrane, likely corresponding to resident membrane-linked proteins, as well as transiting cargo biomolecules. In tomograms collected at 64,000x magnification, these membrane-linked densities could be well resolved, indicating a route towards directly mapping secretory pathway protein organization. To understand how ER/Golgi-resident proteins are arranged within a single, ‘intact’ secretory pathway, we set out to quantitatively examine tomogram Position321, which (serendipitously) captured a full set of ER (cisterna 1), *cis/media/*trans-Golgi (cisternae 2-5), and TGN (cisternae 6-8) compartments, all within the same field of view (**Figure 3a**).

Protein densities within Position321 were surveyed by template matching using the PyTME algorithm^57^. Due to the small size of most proteins within the ER and Golgi (100-200 kDa), we anticipated that direct template searches using known structures would likely perform poorly. Indeed, PyTME matching of Position321 against PDB structures of MGAT1, MGAT5, EXT1/2, NDST1 and C1GALT1 returned implausibly distributed coordinates throughout the entire tomogram, in line with the bias of template matching algorithms towards matching size, rather than specific features^58^ (**Supplemental Figure 5**; **Supplemental Table 2**). Given these technical limitations, we instead turned to examining general biomolecular organization across the secretory pathway, rather than the distributions of specific proteins. By conducting this ‘agnostic’ survey, we reasoned that we could map the distributions of all proteins and protein complexes across Position321, thereby informing upon the fundamental modes of protein organization throughout each compartment of the HeLa secretory pathway.

To conduct a broad survey of protein organization, template matching was again attempted on Position321 using a 6x6x7 nm spheroid, designed to approximate the shape of globular proteins of ∼100-200 kDa in mass^59^ after reconstruction and missing wedge artefacts^60^. As expected, PyTME searches using this model mapped to nearly all densities throughout the tomogram (254,810 picks**; Supplemental Table 2**). To focus on membrane-linked proteins that are principally involved in secretory pathway function, we further discarded all PyTME matches not within 2-9 nm of a segmented ER, Golgi or TGN membrane (13,740 remaining picks). The resulting coordinates were largely localized 6-8 nm from each membrane, and closely matched density projections from *en-face* views, suggesting broadly representative protein sampling across all membranes of Position321 (**Figure 3b**; **Supplemental Figure 6**).

**Figure 3.**
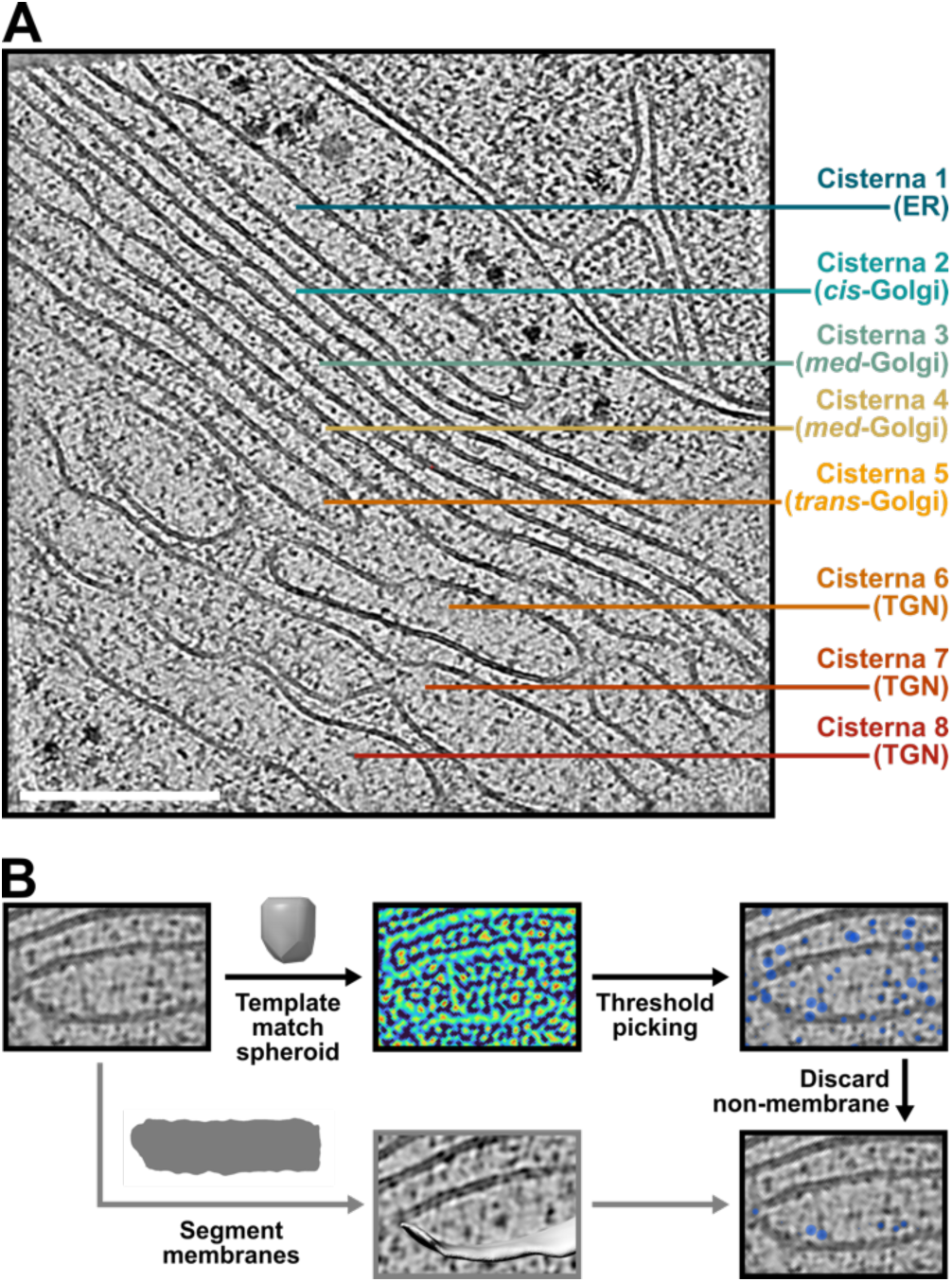
Survey of membrane-linked proteins within a ‘complete’ HeLa secretory pathway. (**A**) Snapshot of tomogram Position321 showing capture of rough-ER, Golgi and TGN cisternae within a single field of view. Distinct membrane linked protein densities are apparent on all membrane surfaces. (**B**) Template matching strategy to survey membrane-linked proteins. Densities are first matched to an ‘agnostic’ spheroidal template, before discarding coordinates not within 2-9 nm of a segmented membrane. Scale bar – 200 nm. See also **Supplemental Table 2**.

### Molecular organization within the HeLa secretory pathway

Our agnostic surveys of Position321 revealed a dense distribution of proteins across each of its ER, Golgi and TGN membranes, with visual inspection suggesting no strict organization across any of the membranes surveyed (**Figure 4**). To quantitatively assess the interplay of these membrane-linked proteins, we first turned to measurements of pairwise distances, revealing a median nearest-neighbor interprotein distance of ∼7.75 nm across Position321. This median nearest-neighbor distance was broadly similar across all ER, Golgi and TGN cisternae surveyed, suggesting that the overall distribution of membrane-linked proteins does not differ significantly between secretory pathway compartments in HeLa cells (**Figure 5a**).

Whilst our survey of interprotein distances did not show substantial differences in nearest-neighbor distributions *between* cisterna (**Figure 5a**), the spread of nearest-neighbor distances *within* each cisterna was substantial. ∼17% (2373/13739) of proteins examined in Position321 possessed a nearest-neighbor within ∼6 nm, less than the diameter of the spheroid used for template-matching, suggesting possible participation in protein-protein interactions. Quantifying the number of proteins involved in such putative interactions revealed that ∼7-10% of proteins within each ER/Golgi cisterna were within close-contact distance (defined as nearest-neighbor <6 nm) (**Figure 5b**). Next, we further assessed the number of proteins involved in dimer, trimer and higher-order interactions using Kluster, a custom-developed program to examine distribution networks. These analyses revealed that HeLa secretory pathway interactions are dominated by small dimeric interactions, with progressively fewer complexes formed as oligomericity increases (**Figure 5c**). As with the general distribution of membrane-linked proteins, these complexes were also stochastically organized, with no strict patterning observed across any ER/Golgi/TGN compartment surveyed (**Figure 4**).

Intriguingly, our examination of protein clusters suggested that certain secretory pathway compartments may be more enriched in higher-order complexes. In particular, the *medial*-Golgi (cisternae 3-4) and late TGN (cisternae 7-8) compartments of Position321 each showed ∼15% of complexes bearing 3 or more subunits, compared to only ∼8% in other cisternae surveyed (**Figure 5c**). Such an increase in higher-order complexes may reflect functional activities, such as the many N-, O- and other types of glycosylation reactions that take place within the *medial*-Golgi^61^. Similarly, increased higher-order complexes in late TGN may reflect trafficking from these compartments, which are ultimately responsible for dispatching biomolecules to their final locations^36^. Further examination of other tomograms will determine whether these changes to protein clustering within *medial-*Golgi and TGN represent a general feature of secretory pathway organization.

**Figure 4.**
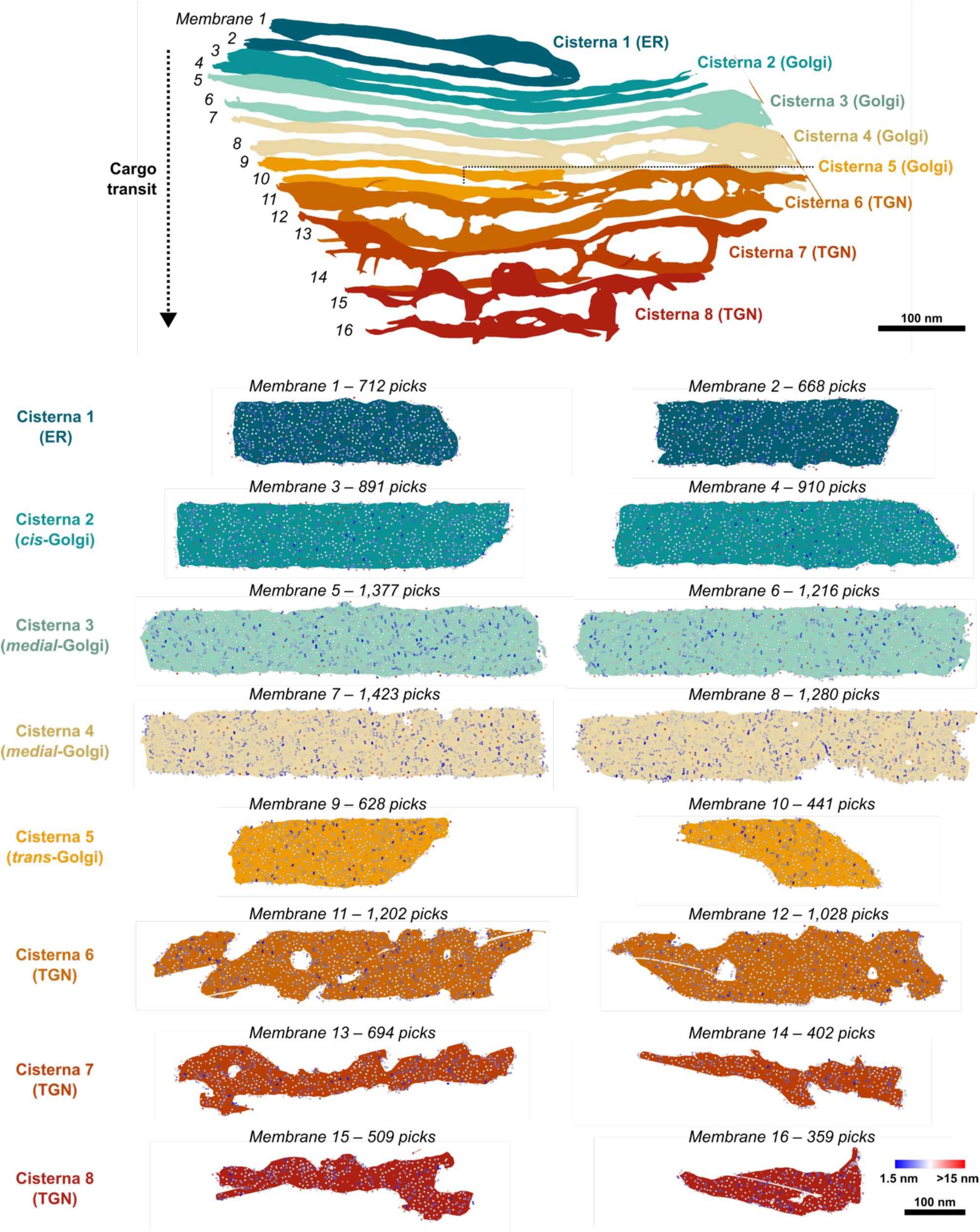
Organization of membrane-bound proteins within the secretory pathway. (**A**) Membrane segmentation of secretory pathway compartments in Position321. (**B**) Agnostic template-matched protein coordinates mapped onto each ER, Golgi and TGN membrane of Position321. Colors of points represent nearest neighbor distances, as described by color bar. Scale bars – 100 nm.

**Figure 5.**
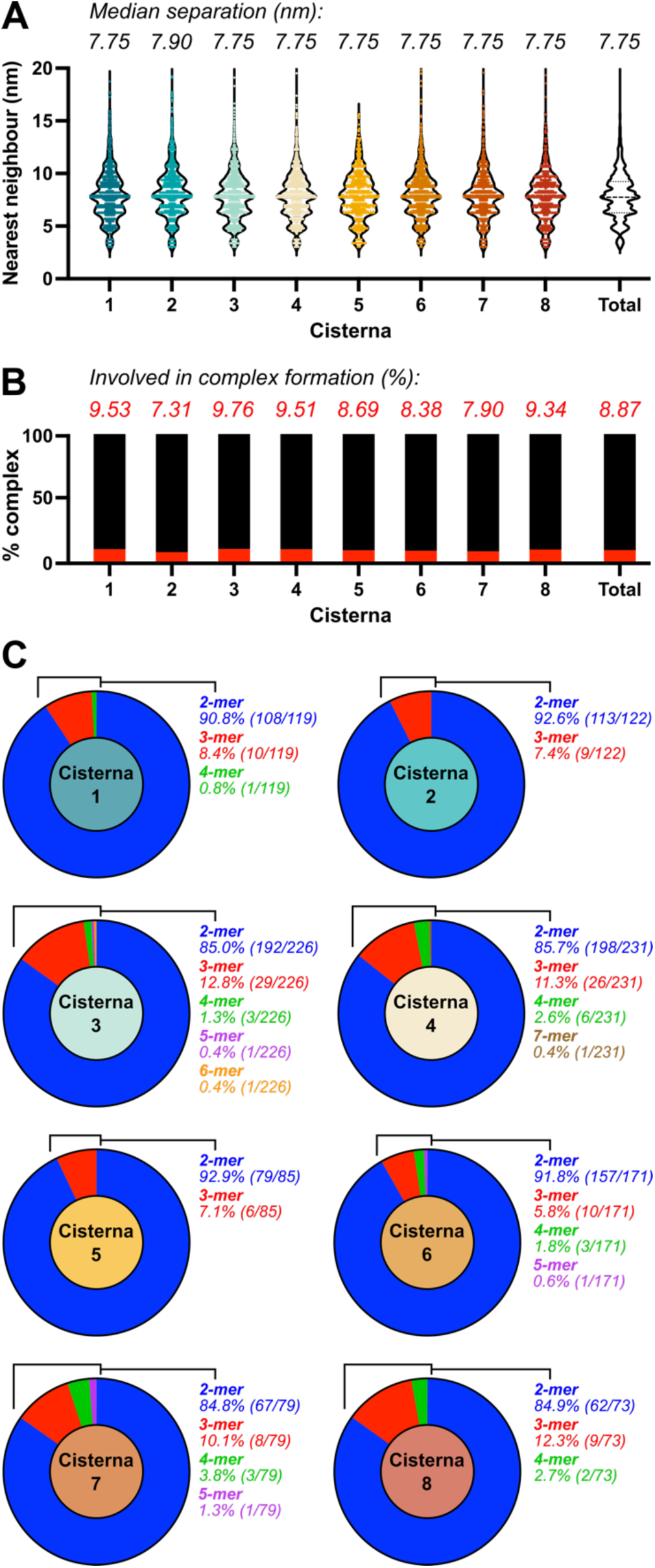
Quantitative survey of secretory pathway protein organization. (**A**) Nearest-neighbor distances for proteins across each secretory pathway cisternae of Position321. (**B**) Percentages of proteins involved in interactions within each cisterna, as defined by a nearest-neighbor distance < 6 nm. (**C**) Proportion of proteins forming dimeric, trimeric, and higher-order contacts within each cisterna.

### Structural features of secretory pathway complexes

Finally, having mapped the organization of membrane-linked proteins across Position321, we turned to direct inspections of identified protein contacts, to examine the molecular basis of their interactions. Selected protein complexes from ER (cisterna 1), *cis*-Golgi (cisterna 2), *medial-*Golgi (cisterna 4), *trans*-Golgi (cisterna 5) and TGN (cisterna 8) are shown in **Figure 6**.

Across Position321, we found that membrane-linked protein complexes were principally comprised of lateral interactions between closely abutting proteins. This lateral mode of interaction appeared consistent regardless of stoichiometry, with higher-order complexes simply presenting more laterally arranged proteins, rather than forming more tightly packed assemblies (*e.g.* **Figure 6** – clusters 4, 12). Furthermore, whilst most complexes we examined formed between proteins originating from the same membrane of a given ER/Golgi/TGN cisterna, a smaller number of contacts also originated from opposing membranes, particularly in regions where cisternae showed narrowing (*e.g.* clusters 7, 10). It is not clear from our data whether such contacts are causally responsible for cisternal narrowing, or whether they may form opportunistically as opposing membranes approach.

Close inspection of Position321 highlighted two distinct modes of protein interaction operating across secretory pathway membranes. In the first mode, two or more proteins formed interactions predominantly *via* their globular head domains (*e.g.* clusters 6, 10, 14), which enabled close interprotein contacts even when their membrane-linking stalks were several nm apart. Functionally, the globular domains of membrane-linked proteins typically bear active sites, such as those involved in ligand binding or catalysis. Because these domains must directly contact their targets for activity, it is plausible that our observed globular domain interactions may correspond, at least in part, to active processing events between resident secretory pathway proteins and transiting cargo biomolecules.

The second mode of interaction we observed comprised pairwise contacts between proteins occurring *via* their stalk domains, which were projected in proximal fashion from ER or Golgi membranes (*e.g.* clusters 3, 18). Compared to interactions formed between globular head domains, these stalk-driven interactions are unlikely directly reflect direct processing events. Instead, such contacts may represent stable complexes between resident ER/Golgi proteins, such as the stalk-driven complexes known to form between enzymes involved in N-glycosylation^62^. By forming such contacts, the functional globular domains proteins would be placed in close proximity, thus increasing the efficiency of any coupled processing compared to isolated, non-interacting proteins.

Taken together, our examination of HeLa secretory pathway proteins reveals a plethora of different interaction modes taking place within ER/Golgi cisternae. Such diversity reflects both the complex dynamic nature of secretory pathway function, as well as our capture of both resident ER/Golgi proteins and transiting biomolecular cargoes.

**Figure 6.**
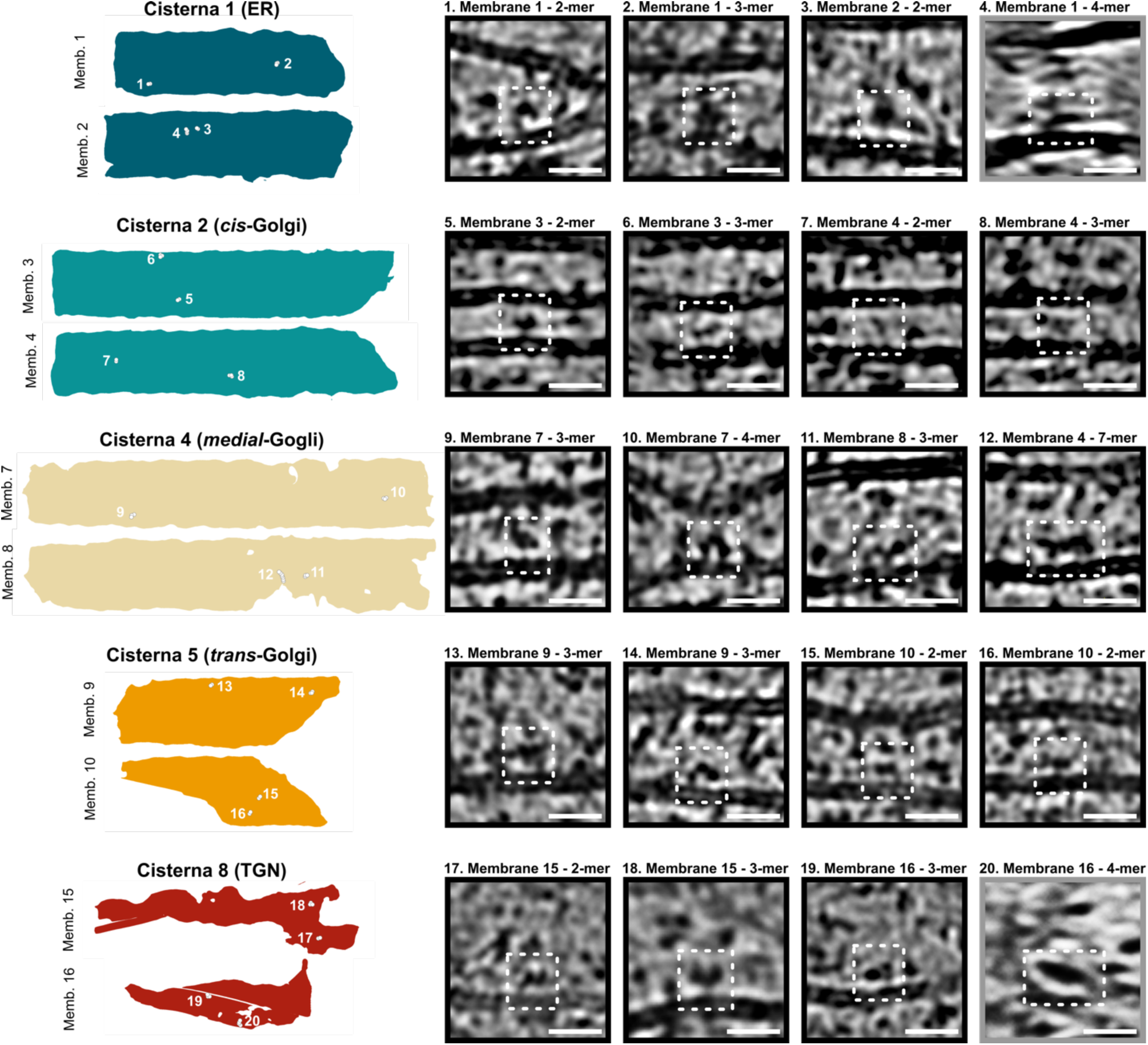
Selected interactions within ER and Golgi cisternal membranes. Complexes between secretory pathway proteins are largely comprised of lateral interactions, occurring both via globular head domains and stalk domains. Most views were taken across the high-resolution tomogram X-Y plane (black borders), Selected complexes were also visualized across the lower-resolution Y-Z plane (grey borders). Scale bars – 20 nm.

## Discussion

The myriad trafficking and processing functions that take place within the eukaryotic secretory system reflect a considerable degree of molecular complexity within its membrane-bound compartments. To date, a comprehensive spatial analysis of protein organization and interaction within the human ER and Golgi has yet to be reported. Here, by exploiting the near-molecular resolutions afforded by cryo-ET, we have mapped both the ultrastructure of the HeLa secretory pathway, and surveyed protein organization across its ER and Golgi membranes. We find that membrane-linked proteins within the HeLa secretory pathway are densely but stochastically distributed, with their interactions dominated by small dimeric contacts occurring from both globular head-domains, and membrane-linking stalk domains. The analyses we have conducted here combine established cryo-ET methods, alongside newly developed tools for analyzing protein distributions and clustering. Our workflows are generalizable, and may be adapted to support further studies of secretory pathway function across both HeLa and other cell-types.

Although technical limitations here have prevented us from identifying specific proteins in our tomograms, the analyses of protein organization we have conducted still enable hypotheses of secretory pathway function to be tested. Notably, for enzymatic processing functions such as glycosylation, a plethora of reported interactions between enzymes in the ER/Golgi^13–23^ has led to speculation of large multi-subunit complexes involved in substrate processing, whose existence would allow multiple interconnected reactions to take place within a single supramolecular assembly^63,64^. Whilst we do observe some higher-order complexes in our tomograms (*e.g.* **Figure 6** – cluster 12), their relative paucity suggests that these are unlikely to be a main driver for intensive processes such as glycosylation. Instead, a predominance of small dimeric protein contacts in our tomograms suggests that enzymatic processing is more likely to occur as a series of reactions between distinct enzyme-substrate pairs, with substrates moving between enzymes to facilitate successive reactions. Notably, our data do not preclude other orthogonal modes of molecular organization that may also influence function, such as compartmentalization of enzymes within sub-regions of cisternae, as has been suggested by super-resolution microscopy^29^. New insights will arise as technical advances enable more accurate annotation of precise proteins/enzymes within tomographic datasets.

Beyond direct analyses of biomolecular organization, our cryo-ET volumes also add to a growing body of data describing secretory pathway structure/function across cell types and species^30–36,38–40^. In this vein, informative comparisons may be drawn between our cryo-ET studies of the HeLa secretory pathway, and recent surveys of *C. reinhardtii* Golgi. A key observation in *C. reinhardtii* tomograms was the presence of arrayed protein densities within their *trans-*Golgi cisternae^38^. Such arrays are conspicuously absent in our tomograms of HeLa Golgi, wherein stochastic intracisternal protein distribution appears to be the norm. Although we do not discount the possibility that array-like protein structures may form transiently within HeLa ER/Golgi (*e.g.* in cell cycle dependent fashion), our data are also suggestive of a fundamentally different mode of molecular organization, highlighting the variability of secretory pathway structure/function that can operate between cell types and species. Further studies of ER and Golgi at the molecular level are needed to resolve fundamental principles that operate in common between species, thereby bringing us closer towards generalized models for secretory pathway function.

## Methods

### Cell culture

HeLa cells were cultured in Dulbecco’s modified eagle medium (DMEM) [Gibco], supplemented with 10% v/v fetal bovine serum (FBS) [Gibco], 1% v/v penicillin/streptomycin [Gibco] and 1% v/v non-essential amino acids [Gibco]. Cells were maintained below 90% confluency and 30 culture passages.

### Fluorescent protein cloning

Donor plasmids encoding for full length EXT1, EXT2, NDST1, MGAT1, MGAT5 and C1GALT1 were obtained from the DNASU repository [Arizona State University]. For full length GFP tagged proteins, cDNA fragments encoding for genes of interest were excised by PCR (Q5 DNA polymerase [NEB]), and cloned by Infusion [Takara] in frame into a pOPINE-3c-eGFP vector^65^, previously linearized at the NcoI + PmeI restriction sites. The resulting constructs encoded for full length proteins, each fused to a C-terminal eGFP fluorophore.

For GFP11-tagged EXT2, full length EXT2 was first appended to a DNA fragment encoding for N-terminal GFP11, before PCR amplification to create the tagged cDNA product. GFP11-EXT2 was cloned by Infusion into the pOPINE vector, previously linearized at the NcoI + PmeI restriction sites.

For GFP(1-10), a gene encoding for the 1^st^ 10 β-sheets of GFP was cloned by Infusion into the pHR-AIO vector [Takara], linearized at the EcoRI and XhoI sites. The resulting plasmid contained GFP(1-10) integrated behind the Tet/Dox-responsive TRE3GS promoter.

All constructs were verified by Nanopore sequencing [Source Biosciences] before further use.

### Stable split-GFP HeLa cell line cloning

HeLa cells containing stable integration of the GFP(1-10) cassette were produced using lentiviral transduction, using lentivirus produced in the Lenti-X system [Takara]. Briefly, Lenti-X 293T cells in grown in DMEM, 10% v/v FBS and 1% v/v penicillin/streptomycin, were transfected with pHR-TRE3GS-GFP(1-10) using Lenti-X packaging single shots, which contain Xfect transfection reagent pre-mixed with VSV-G-pseudotyped packaging plasmids. Following 24 h incubation, lentivirus containing media was harvested, filtered through a 0.45 µm PES filter, then snap frozen in single use aliquots for further use. Viral titres were determined using the Lenti-X GoStix Plus assay [Takara].

For transduction, HeLa cells in a 6-well plate were treated with a solution of DMEM containing polybrene (4 µg/mL) and lentivirus at 1:10 dilution. Transduced cells were incubated for 24 h, before changing media for fresh DMEM, and further incubation for 48 h. Transfected cells were progressed to monoclonal selection in 96-well plate format, with single cell sorted conducted by FACS. Doxycycline induction of GFP(1-10) in monoclonal cells was examined by western blotting using HRP-conjugated anti-GFP [Invitrogen A10260], before expanding verified clones for further studies.

### Fluorescent gene expression

Plasmids encoding for full length GFP-tagged proteins of interest were transfected into HeLa cells using FuGene HD following manufacturers’ instructions, at a typical w/v ratio of 2 µg DNA: 5 µL FuGene reagent. Cell media was exchanged for fresh media 24 h following transfection. Typically, fluorescence was visible 24-48 h following transfection. For fluorescence imaging, cells were further stained with Hoechst 33258 [Invitrogen], before visualization using a Leica SP8 confocal microscope.

For split-GFP fluorescence, HeLa-GFP(1-10) cells transfected with EXT1 + GFP11-EXT2 were treated with 1 µg/mL Doxycycline 24 h after transfection. Visible fluorescence signal typically appeared 24 h following Doxycycline treatment. Fluorescence imaging was conducted as described above.

### BODIPY-ceramide staining

HeLa cells were treated with DMEM media supplemented with BODIPY FL C5-Ceramide [Invitrogen] at varying v/v ratios and incubation times (**Figure 1**). For fluorescence imaging, cells were further stained with Hoechst 33258, and imaged by confocal microscopy as described above.

### HeLa cell grid seeding and vitrification

Trypsinized HeLa cells were seeded onto Quantifoil 200 R2/2 Au grids [Jena Bioscience], glow discharged in a GloQube Plus glow discharger [Quorum Technologies] at 30 mA negative current for 60 s. Glow discharged grids were placed into a tissue culture dish, and seeded with cells at a density of 30,000 cells/cm^2^, resulting in an approximate distribution of 1 cell per EM grid square. For cells labelled by fluorescent protein transfection, GFP-positive transfected cells were first enriched using a BioRad S3e cell sorter, before seeding this sorted population onto grids. For ceramide labelling, grids were seeded with wild-type HeLa cells and incubated for 24 h, before treating with fresh DMEM containing BODIPY FL C5-Ceramide as described above.

Cells were typically vitrified for imaging 24 h after seeding onto grids. Quantifoil grids carrying cells were first briefly passed through a solution of DMEM media supplemented with 10% glycerol [Merck] (typically 5-10 s contact time). Grids were then immediately processed using a Vitrobot Mark IV [Thermo Fisher Scientific (TFS)] set to a chamber temperature of 20 °C, with 100% relative humidity. Grids were backside blotted by attaching a Teflon sheet to the Vitrobot front-side pad. Blots were carried out for 10-15 s using a nominal blot force of -10, before plunge freezing into liquid ethane. Grids were clipped into Autogrid rings [TFS] before loading into microscopes for screening, milling, and data collection.

### Cryo-FIB milling

Vitrified HeLa cells were processed using a dual-beam plasma focused ion beam scanning electron microscope (pFIB/SEM; TFS Arctis). First, grids were inspected by SEM imaging to determine adequate seeding of cells, and other quality metrics (grid damage *etc*). Selected grids were then further screened using the integrated fluorescence microscope of the Arctis, to identify regions of interest for milling. Automated pFIB milling was conducted using the AutoTEM 4 software [TFS] and Argon plasma, as previously described^66^. Briefly, rough milling steps at 2.0 nA, 0.74 nA and 0.2 nA were first used to remove material above and below the intended lamella position. This left a lamella of ∼700 nm thick, which was further thinned and “polished” using 60 pA and 20 pA Argon plasma to a final nominal lamella thickness of < 200 nm. Final fluorescence screening was conducted on milled lamella to confirm regions of interest remained within the milled volume.

### TEM data acquisition and reconstruction

Tilt series were collected on milled lamellae using a Titan Krios [TFS] transmission electron microscope (TEM) equipped with a Falcon 4i camera and a Selectris energy filter. Dose-symmetric tilt-series were collected using Tomo 5 software [TFS] at magnifications of 42000x and 64000x (nominal pixel sizes 3.05 Å and 1.90 Å respectively). Tilt series data were collected in electron counting mode from +60° to -60° in 2.5° increments, accounting for the pre-tilt of the lamellae. Total dose of the tilt series was 120–150 e^−^/Å^2^.

For reconstruction, Warp 1.0.9^67^ was used to conduct initial gain and motion correction (10 frame groups per tilt) and contrast transfer function (CTF) correction, and for creating corrected tilt-series stacks. Tilt-series were also manually inspected to remove any clearly erroneous tilts (*i.e.* due to grid-bar incursion, ice-contamination, or excessive shifts from poor tracking). Corrected tilt-series were aligned and reconstructed at binning factors 4 or 8 using AreTomo 1.3.4^68^. Further post-processing was conducted using IsoNet^69^. Tomograms were first subjected to CTF deconvolution (SNR fall-off = 0.7, deconv_strength = 0.7-1.0 depending on dataset). Correction was then carried out using a custom IsoNet model, trained on 20 high-quality volumes collected from the same session as our main tomography dataset. IsoNet model training was conducted using a cube size of 32, and a crop size of 128.

### Tomogram segmentation

All initial tomogram segmentations were performed with MemBrain-seg^70^, using isonet filtered bin4 tomograms as inputs. For each tomogram, MemBrain-seg outputs were subsequently proof-read manually in napari^71^, using the segmentation toolbox plugin, and rendered in AmiraX (2023.1.1).

### Secretory pathway ultrastructural metrics

To assess distances between Golgi membranes, tomogram segmentations were first filtered to remove any ‘small’, disconnected volumes below 50,000 voxel^3^ in size, leaving only volumes corresponding to *bona fide* membranes. For each remaining membrane, surfaces were reapproximated using a dense triangular mesh, with the coordinates of vertices and faces determined using the marching cubes algorithm^72^, implemented in scikit-image^73^. For Position190, this process resulted in 10 meshes (corresponding to 10 membranes), with sizes ranging from ∼142,000 to ∼734,000 points.

For each pair of membranes, we further calculated the distances between every face point on a ‘reference’ membrane to its nearest face point on a ‘search target’ membrane. ‘Reference’ was defined as the smaller membrane within a given membrane pair, and ‘search target’ was defined as the larger membrane. This created a continuous ‘heatmap’ of differences between membrane pairs. Due the computational difficulty of finding closest points using a traditional full pairwise distance matrix, we adopted the k-dimensional tree (“k-d tree”) algorithm^74^ from the scipy library^75^, which allows for efficient evaluation of the closest distances between two ensembles of coordinates. The final set of resulting distances in voxels were converted to nanometers by multiplying against the voxel spacing (1.22 nm/vx). Plots of distance graphs and modal peak calculations were carried out in Prism 11 [Graphpad].

### Template matching and particle subtraction

Template matching of tomograms was conducted using the Python Template Matching Engine (PyTME) version 0.3.3^57^. For protein structure based picking, coordinate models were obtained from the PDB, and converted into 3D volumes using ChimeraX^76^ molmap function. The 0.6x0.6x0.7 nm spheroidal template for picking was generated in ChimeraX. Template masks were generated using the PyTME preprocessor interface. Tomogram masks were generated using SLABIFY. All template matching searches were conducted on bin8 tomograms, using a PyTME picking threshold of 0.55.

To obtain particle distributions focused around membranes, a Python package named Korpuskulum was developed. Korpuskulum uses a volumetric binary mask of a segmented membrane and coordinates of the tomogram particles of interest as input, before evaluating the particle set using the following steps.

First, Korpuskulum internally trims the input segmentation mask (which has the same size as the original tomogram) to the tightest bounding box around the segmented membrane to reduce the search space. This is essential to reduce the computational overhead of the following steps. For each of the slices in the trimmed Z-stack, Korpuskulum calculates the full pairwise distance matrix between the ensemble of particles and the ‘membrane’ voxels in the slice. Hence for *N* particles and *M* membrane voxels, this step would produce an *N* × *M* matrix. The nearest membrane voxel for a particle is defined as the one with the minimum distance in this matrix: for *N* particles, this produces a 1-D list with length *N*. Voxel coordinates corresponding to those particles in the list are also extracted and stored in an *N* × 3 array. This process is repeated for all slices (S), producing a final *N* × *S* × 3 matrix, with full information of the closest membrane voxel per slice. The global minima are extracted by taking the voxels with the lowest overall distance. Mathematically this process can be expressed as

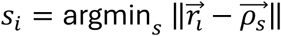

where *s_i_* denotes the voxel index globally closest to the particle *i*, 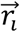 is the coordinates of particle *i*, and 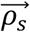 is the coordinates of the membrane voxel indexed *s*. The displacement vector 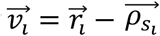 conveniently depicts the orientation of particle *i* with respect to its closest membrane voxel *s_i_*, with the sign of its *z* component signifying the side of the membrane on which the particle is positioned. The norms of the displacement vectors, intuitively denoting the absolute shortest distance between the particles and the membrane, are aggregated and plotted as histograms showing the distribution of particle-membrane separations.

### Particle clustering analysis

Particle clustering analysis was conducted using a newly developed tool named Kluster. Kluster uses the particle output from Korpuskulum to gauge the particle clustering behaviours using two levels of calculations.

The first level of evaluation looks for micro-clusters, i.e. “polymerised” particles, in the ensemble. Kluster first calculates the pairwise distance matrix within the ensemble, then extracts a neighbour list for particle pairs with separation within a user-defined distance threshold, e.g. 4 nm. The neighbour list is then used to build a graph using the NetworkX package^77^ to visualise and quantify the size and quantity of micro-clusters. Kluster does not impose any shape restrictions on the micro-clusters focusing on the requirement for adjacent particles to be within the distance threshold.

Kluster also performs evaluation in a more macroscopic level to gauge the distribution of the clusters. In this calculation, Kluster first calculates the geometric centroids of all the micro-clusters in the system determined in the previous level and including those with a cluster population of 1 (i.e. isolated particles), then evaluates the pairwise distance matrix of the clusters. Using another user-defined parameter of search radius, e.g. 50 nm, Kluster aggregates, for each cluster (“reference”), all the positions of other clusters within the search radius and calculates the ensemble centroid. This is then subtracted from the reference centroid to obtain the displacement which serves as a metric of distribution imbalances. All the moduli of the displacement vectors are averaged to form the overall displacement. A large value for the overall displacement may indicate a biased distribution of particles.

## Acknowledgements

The Rosalind Franklin Institute is a UKRI EPSRC funded institute. LW and CMW acknowledge support from the Wellcome Trust (Sir Henry Dale fellowship to LW - 218579/Z/19/Z). MCD and NByY acknowledge support from the Wellcome Trust through the Electrifying Life Science grant 220526/Z/20/Z

## Author contributions

LW and CMW conceived and developed the project. CMW carried out all cloning, transfections, labelling, and cryo-ET data collections. MAL conducted confocal microscopy. CMW and LW conducted cryo-ET data processing, reconstructions, segmentations and template matching. NByY wrote software and conducted all distance and clustering analyses on coordinate datasets, with guidance from MCD. LW wrote the manuscript with input from all authors.

