## Supplemental Information for "Molecular organization of the eukaryotic secretory pathway"

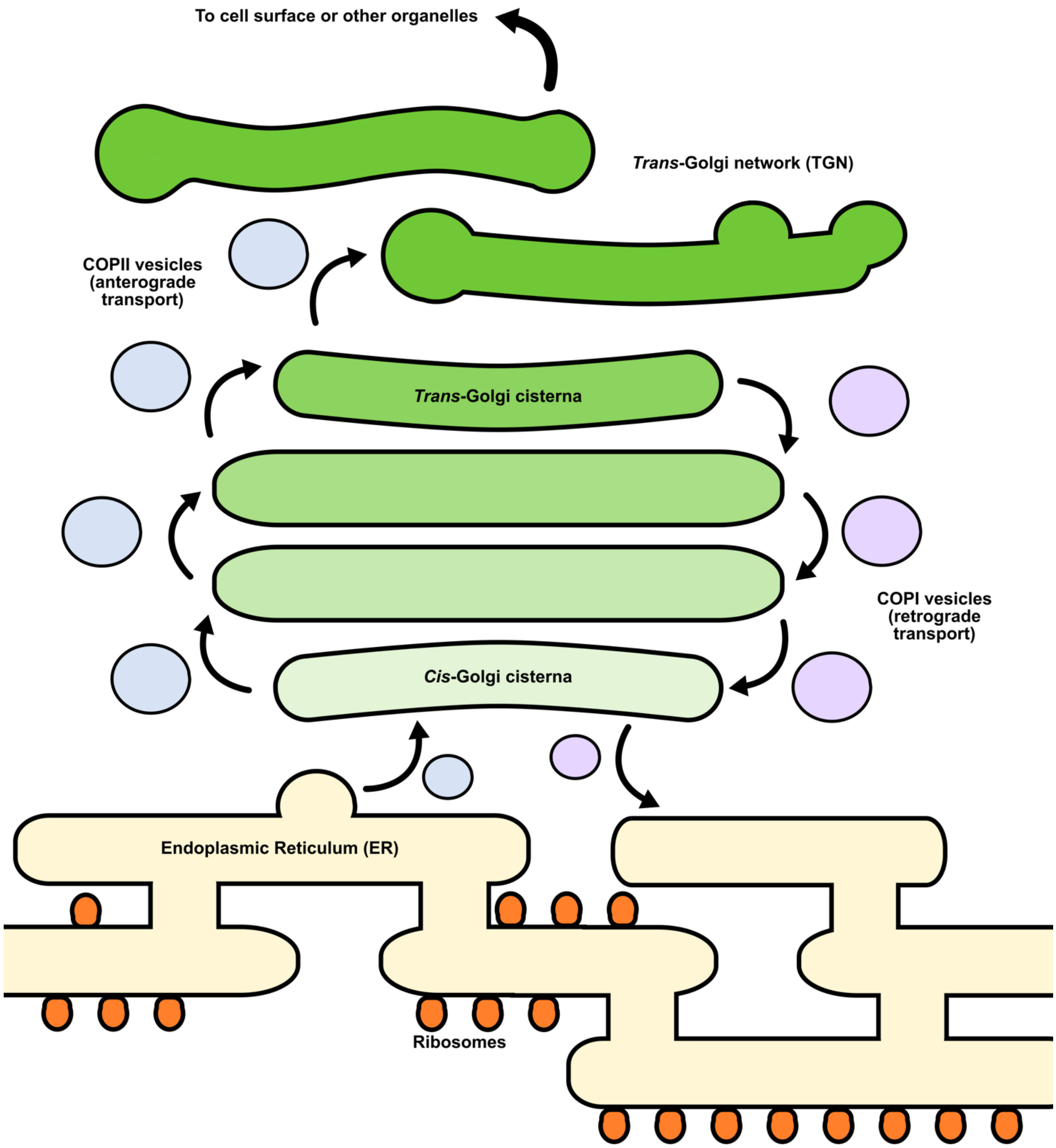


***Supplemental Figure 1*** *Schematic representation of eukaryotic secretory pathway trafficking. Cargo biomolecules are first processed in the endoplasmic reticulum (ER), before being moved to the cis-Golgi cisterna via COPII vesicle transporters. Cargoes transit from cis- to trans-through the Golgi stack, before exiting via the trans-Golgi network (TGN) to further destinations.*

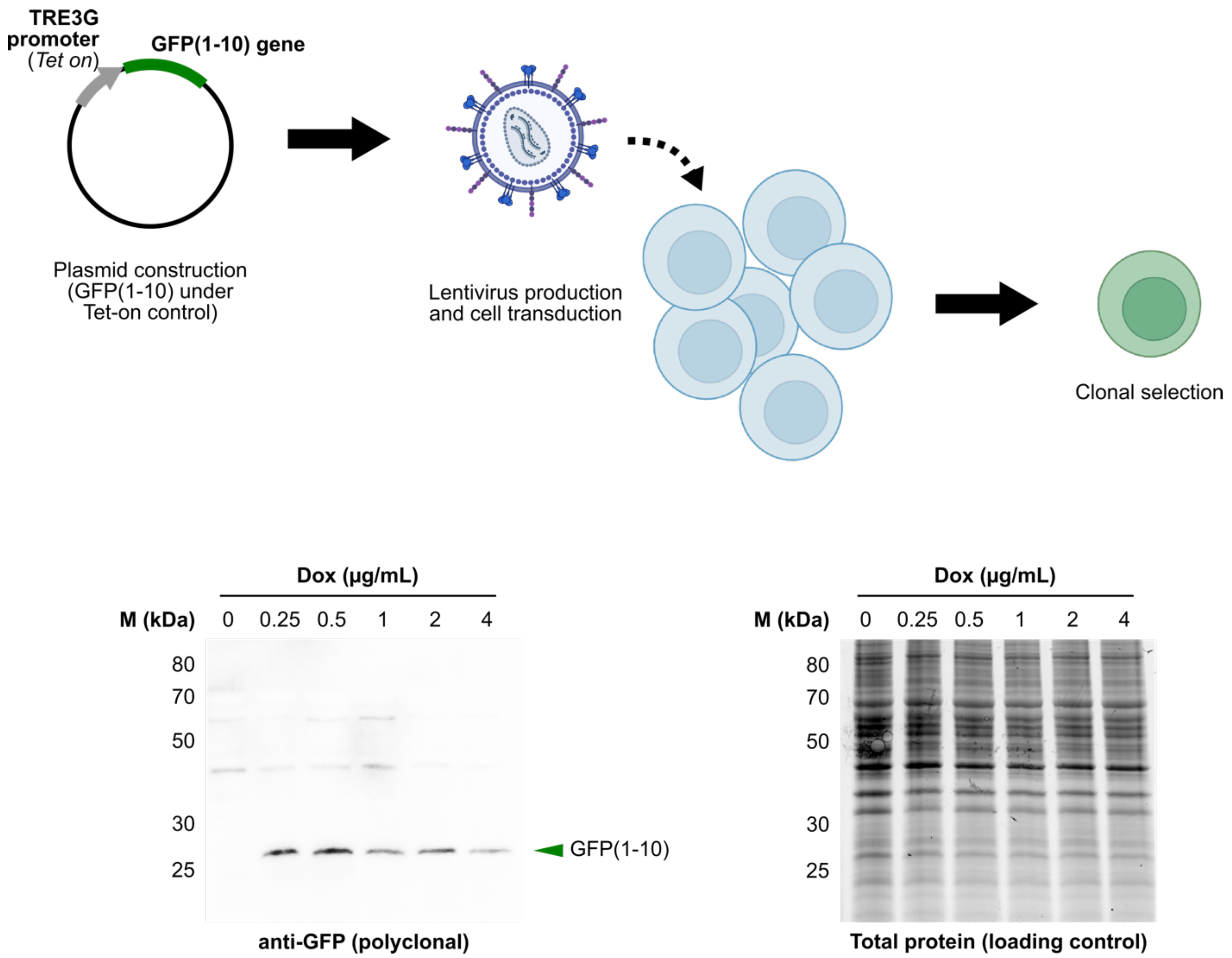


***Supplemental Figure 2*** *Construction of clonal Tet-inducible GFP(1-10) HeLa line by lentiviral transduction, and confirmation of GFP(1-10) expression by western blot.*

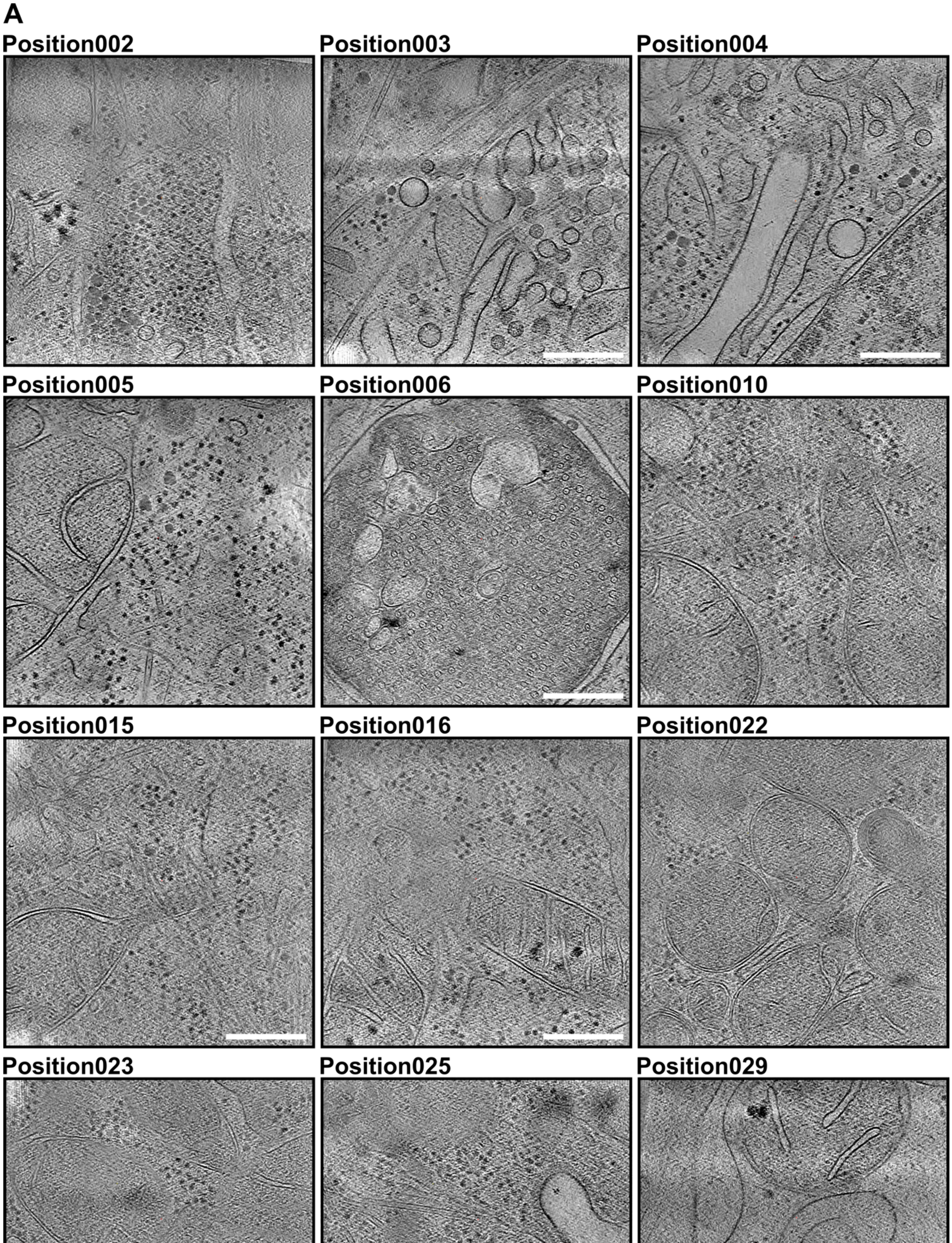
A
Position002
Position003
Position004
Position005
Position006
Position010
Position015
Position016
Position022
Position023
Position025
Position029

**A** cont.

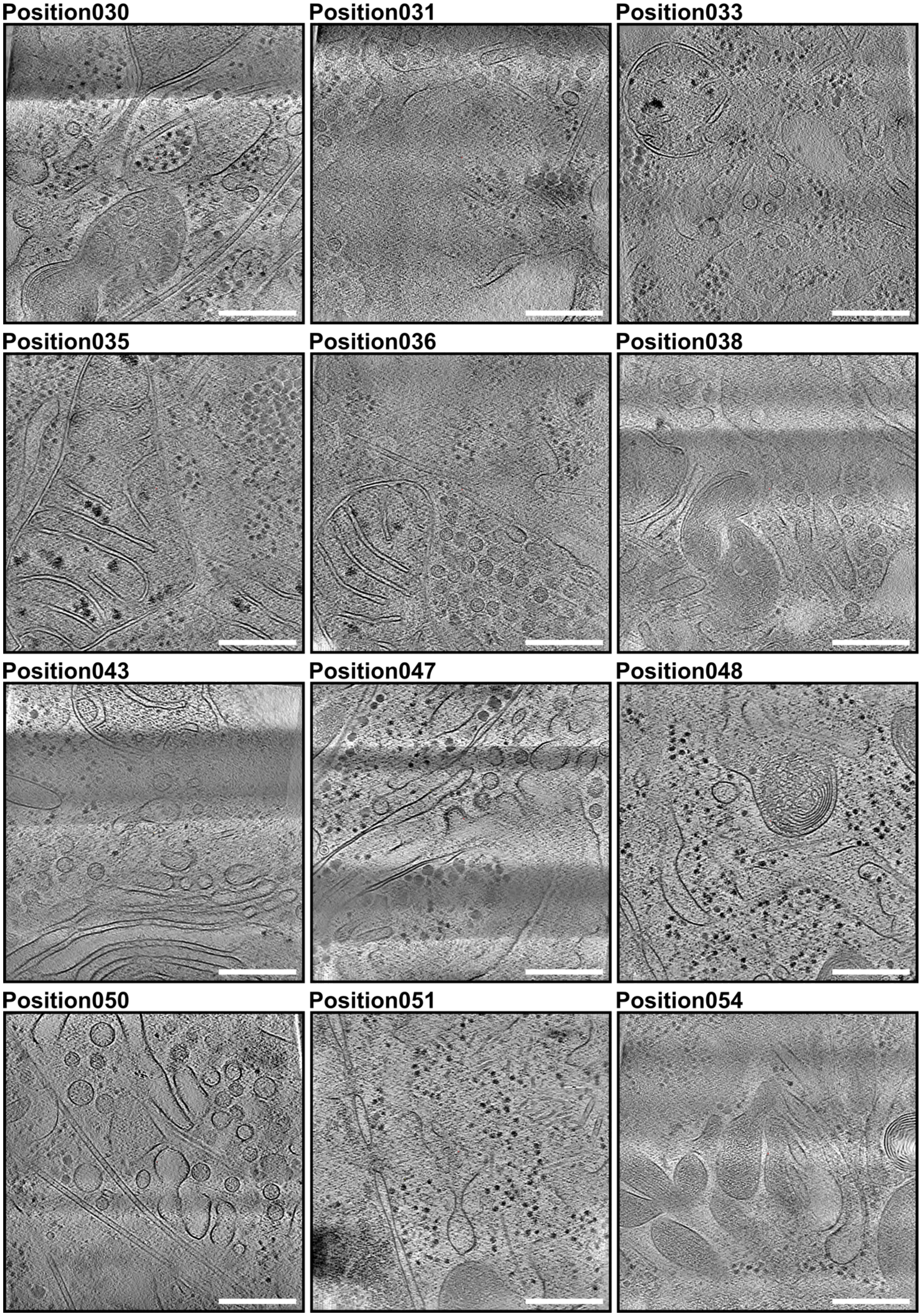
Position030
Position031
Position033
Position035
Position036
Position038
Position043
Position047
Position048
Position050
Position051
Position054

**A** cont.

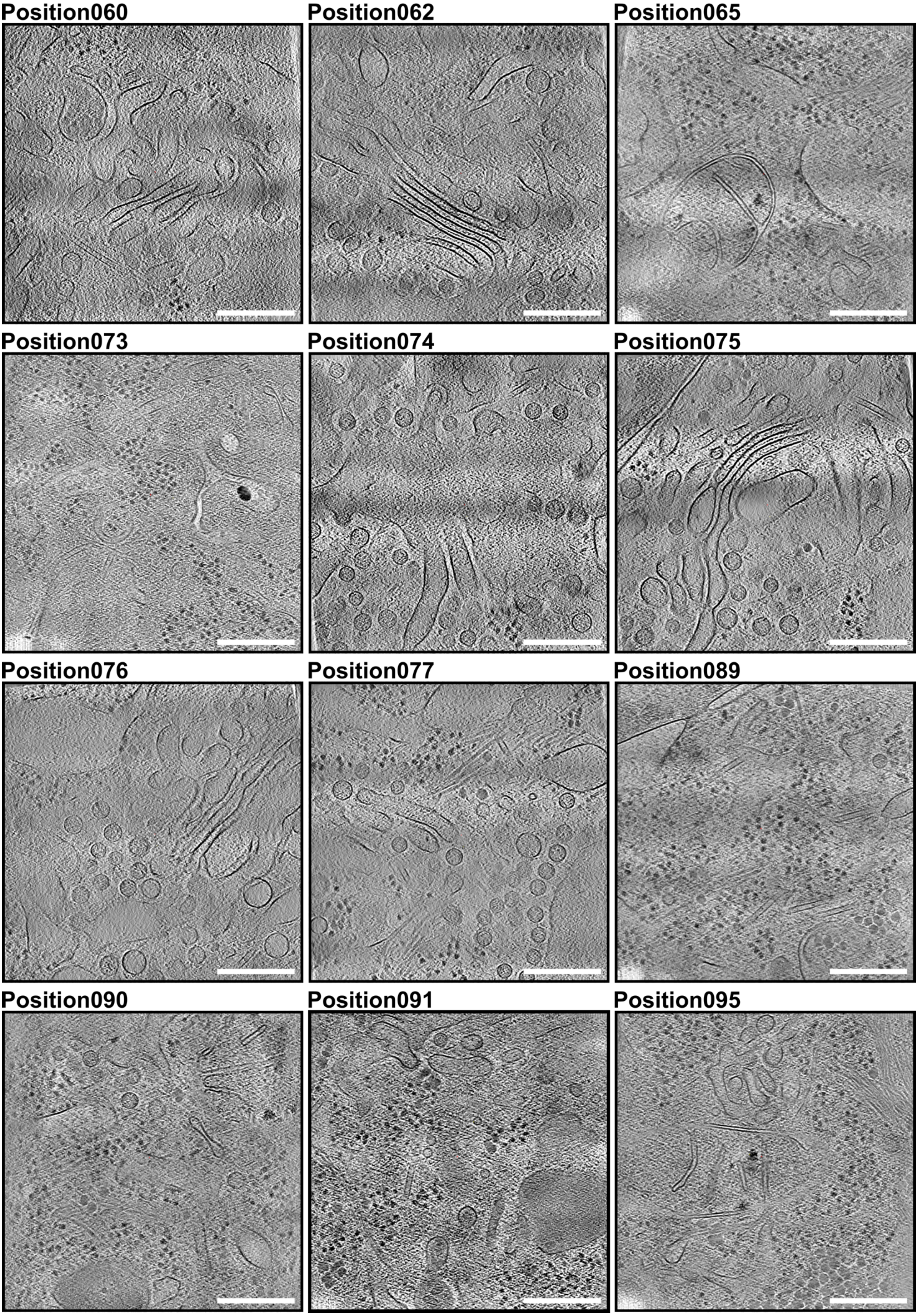
Position060
Position062
Position065
Position073
Position074
Position075
Position076
Position077
Position089
Position090
Position091
Position095

A cont.

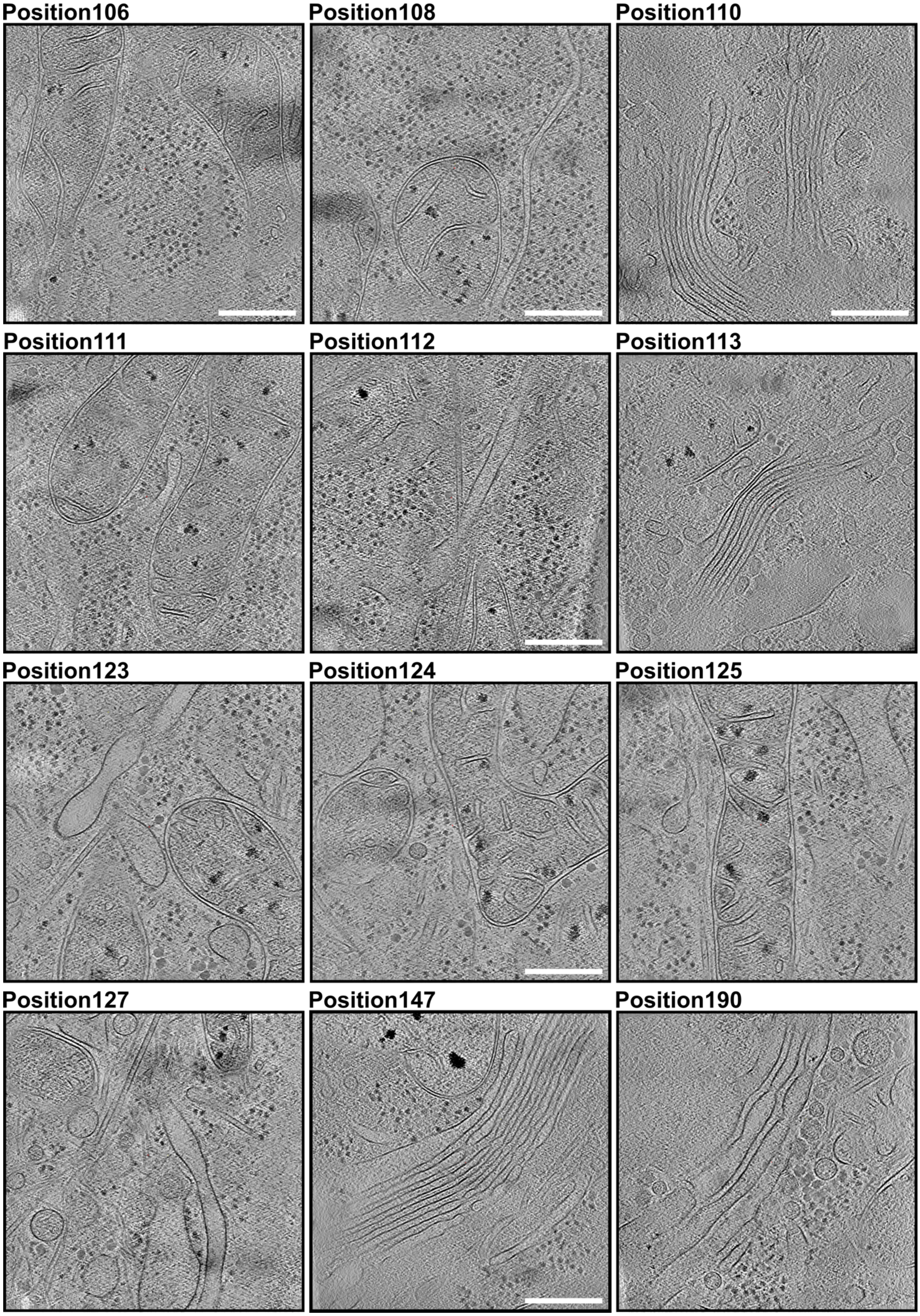
Position106
Position108
Position110
Position111
Position112
Position113
Position123
Position124
Position125
Position127
Position147
Position190

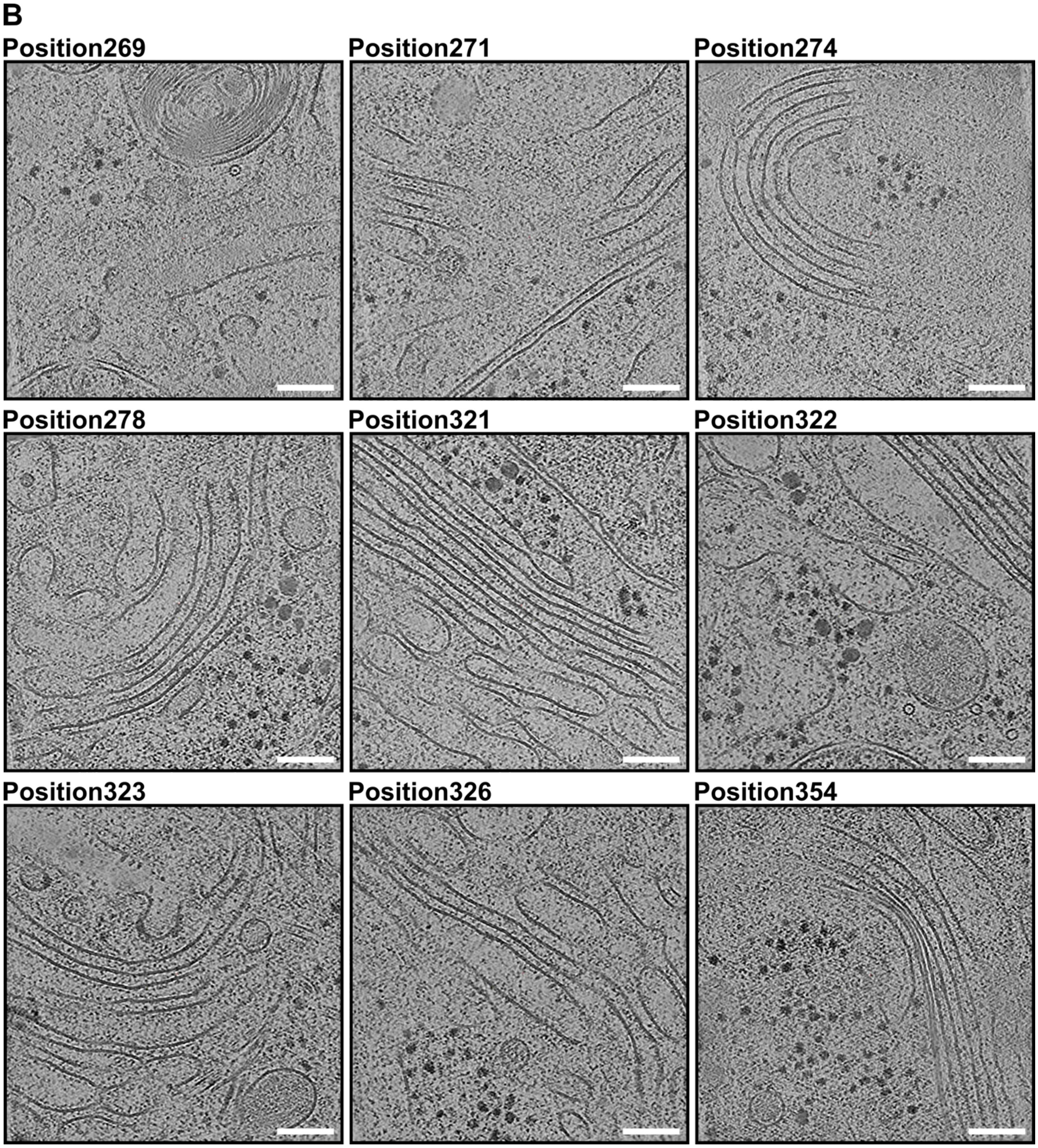


***Supplemental Figure 3*** *(**A**) Overview of selected secretory pathway tomograms collected at 42,000x magnification (nominally 3.05 Å/pixel), and at (**B**) 64,000x magnification (1.90 Å/pixel). Scale bars in A – 200 nm. Scale bars in B – 100 nm.*

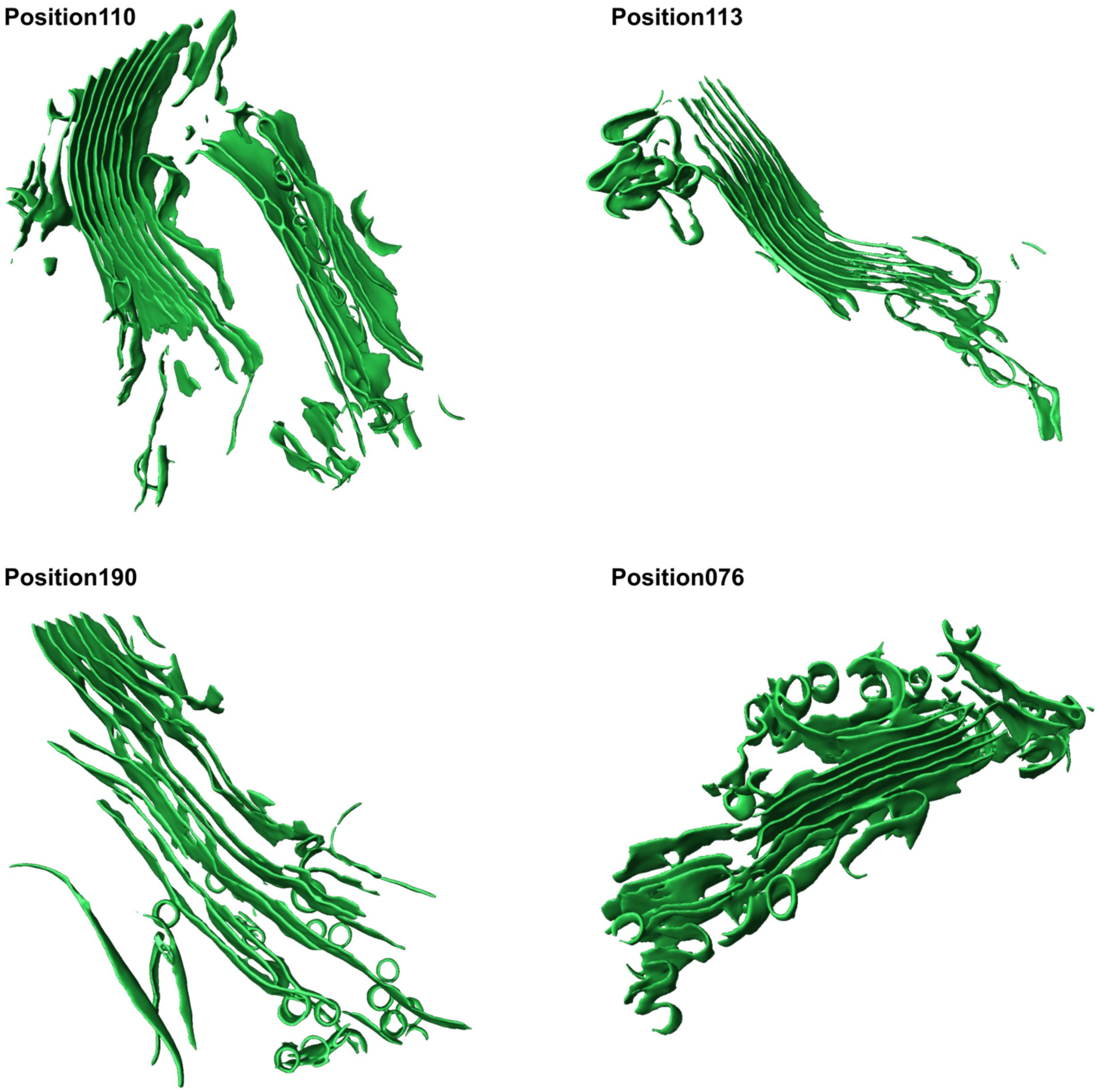


***Supplemental Figure 4*** *Segmented Golgi stacks from tomograms Position110, Position113, Position190, Position075. See also main text* ***Figure 2****.*

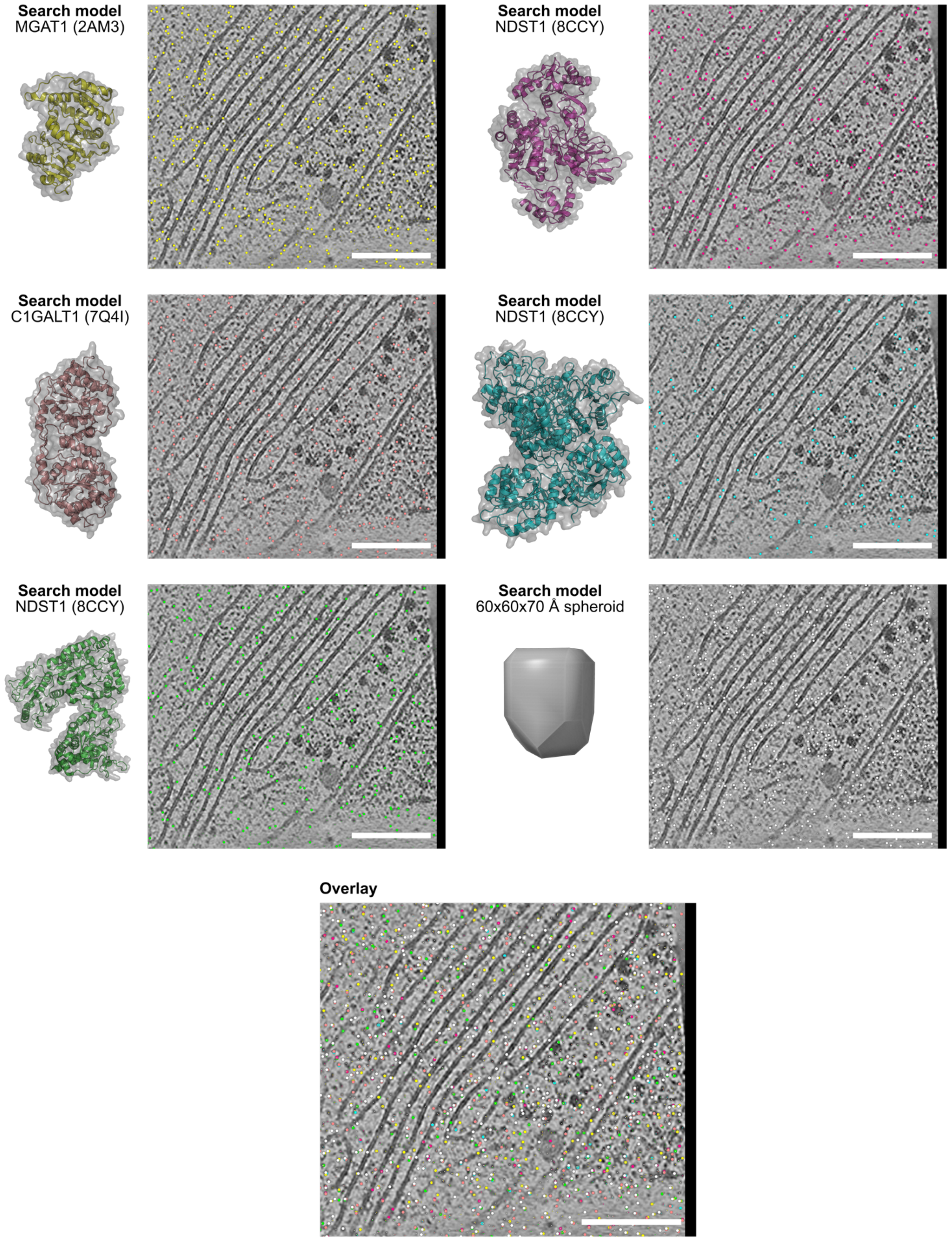


***Supplemental Figure 5*** *Summary of PyTME picking on Position321 using known protein structure templates, shown against a single tomogram slice. Picks are implausibly distributed across the entire tomogram, highlighting poor performance using these templates. Also shown are the picks arising from the spheroidal model used for 'agnostic' template matching. Scale bar – 200 nm.*

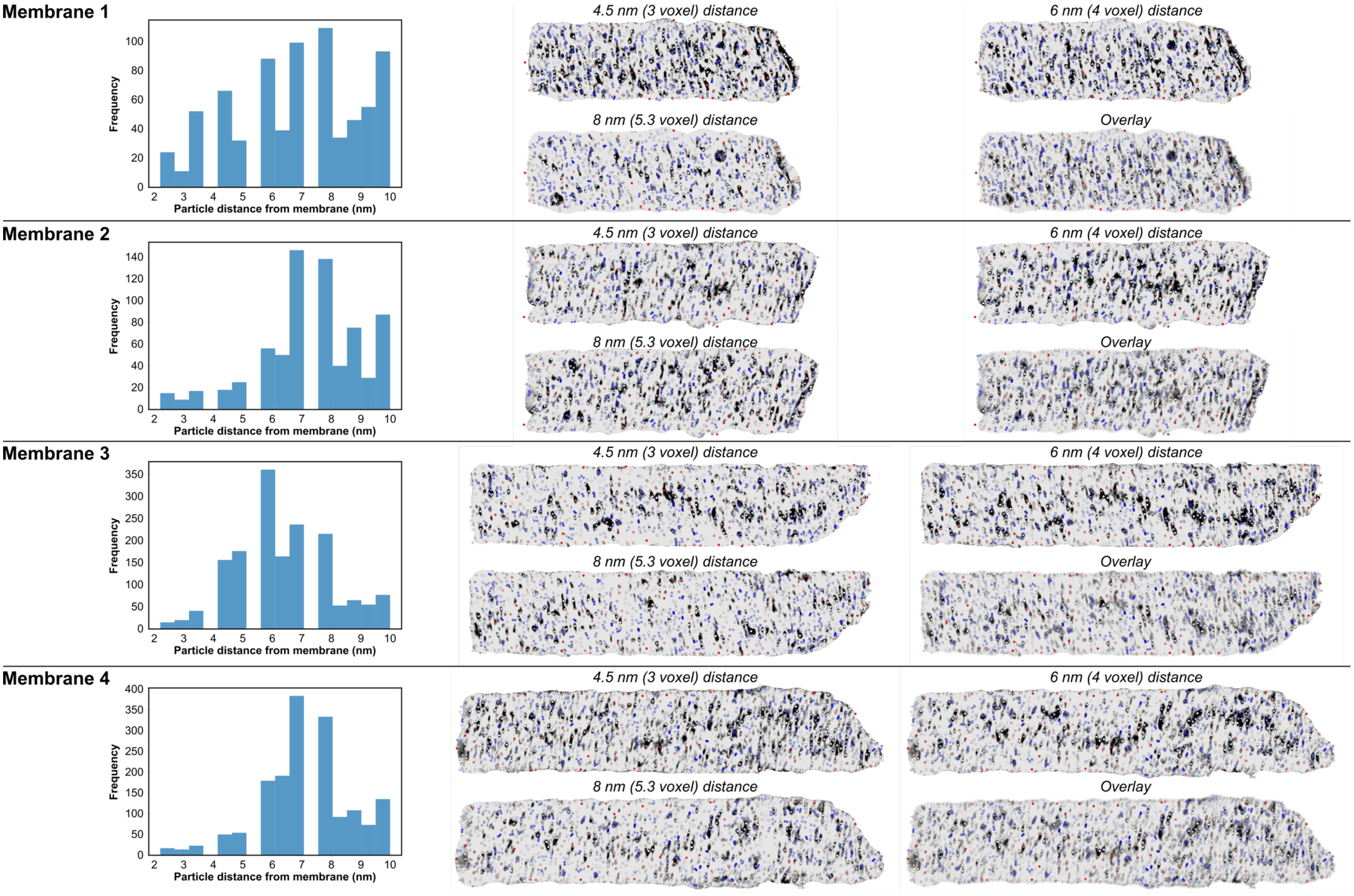

Membrane 1
Frequency
Particle distance from membrane (nm)
4.5 nm (3 voxel) distance
6 nm (4 voxel) distance
8 nm (5.3 voxel) distance
Overlay
Membrane 2
Frequency
Particle distance from membrane (nm)
4.5 nm (3 voxel) distance
6 nm (4 voxel) distance
8 nm (5.3 voxel) distance
Overlay
Membrane 3
Frequency
Particle distance from membrane (nm)
4.5 nm (3 voxel) distance
6 nm (4 voxel) distance
8 nm (5.3 voxel) distance
Overlay
Membrane 4
Frequency
Particle distance from membrane (nm)
4.5 nm (3 voxel) distance
6 nm (4 voxel) distance
8 nm (5.3 voxel) distance
Overlay

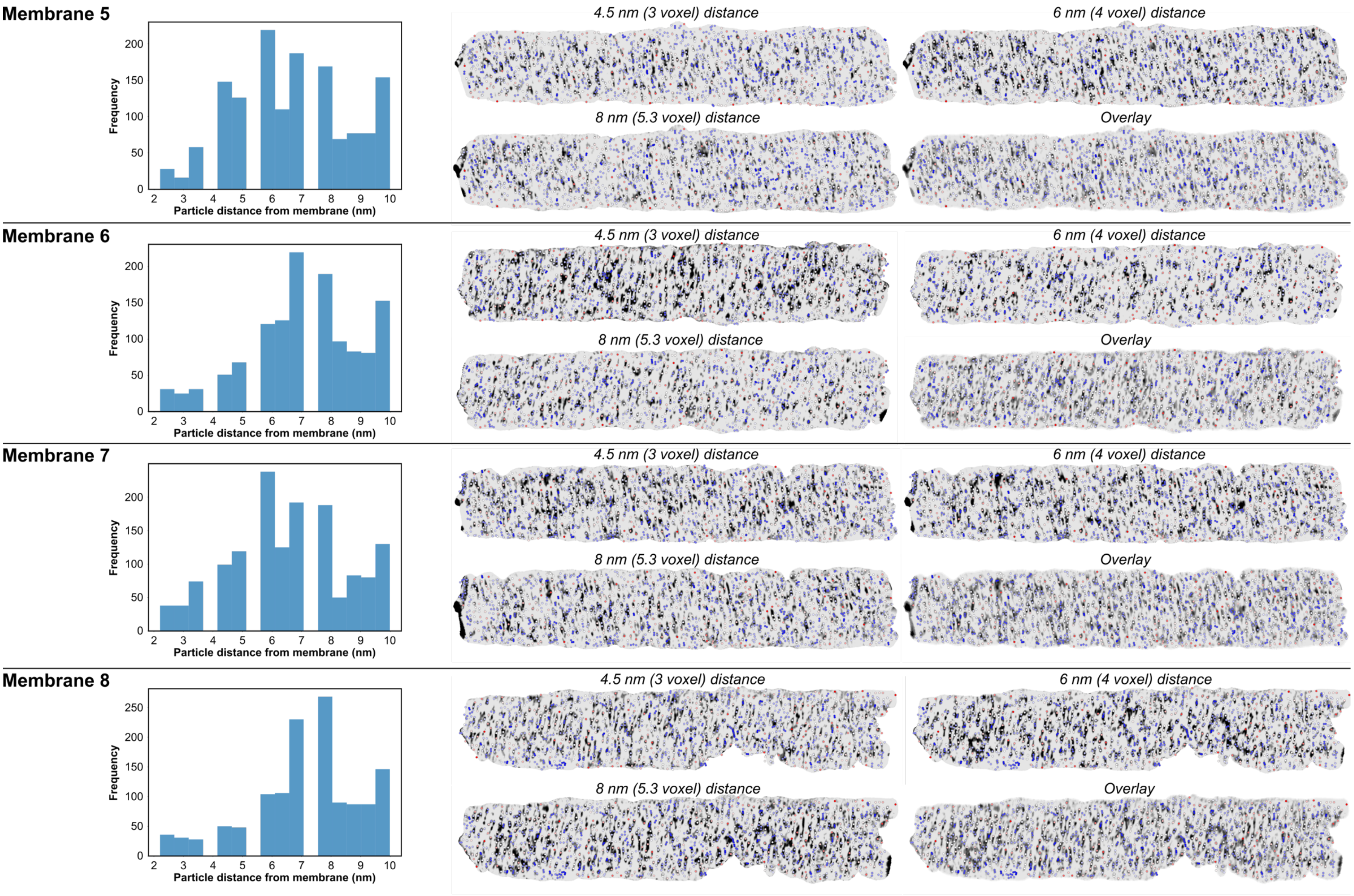
Membrane 5
Frequency
Particle distance from membrane (nm)
4.5 nm (3 voxel) distance
6 nm (4 voxel) distance
8 nm (5.3 voxel) distance
Overlay
Membrane 6
Frequency
Particle distance from membrane (nm)
4.5 nm (3 voxel) distance
6 nm (4 voxel) distance
8 nm (5.3 voxel) distance
Overlay
Membrane 7
Frequency
Particle distance from membrane (nm)
4.5 nm (3 voxel) distance
6 nm (4 voxel) distance
8 nm (5.3 voxel) distance
Overlay
Membrane 8
Frequency
Particle distance from membrane (nm)
4.5 nm (3 voxel) distance
6 nm (4 voxel) distance
8 nm (5.3 voxel) distance
Overlay

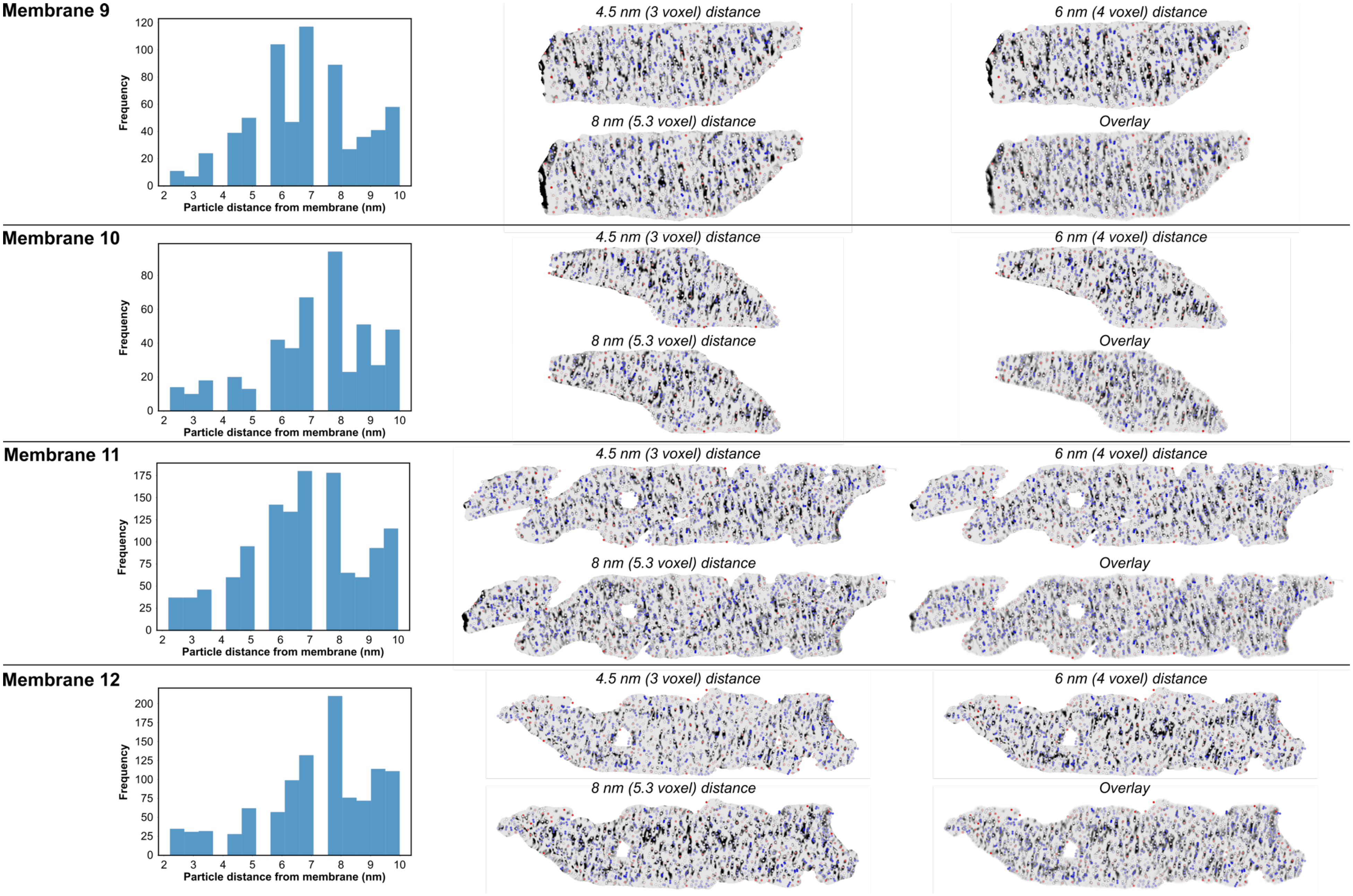
Membrane 9
Frequency
Particle distance from membrane (nm)
4.5 nm (3 voxel) distance
6 nm (4 voxel) distance
8 nm (5.3 voxel) distance
Overlay
Membrane 10
Frequency
Particle distance from membrane (nm)
4.5 nm (3 voxel) distance
6 nm (4 voxel) distance
8 nm (5.3 voxel) distance
Overlay
Membrane 11
Frequency
Particle distance from membrane (nm)
4.5 nm (3 voxel) distance
6 nm (4 voxel) distance
8 nm (5.3 voxel) distance
Overlay
Membrane 12
Frequency
Particle distance from membrane (nm)
4.5 nm (3 voxel) distance
6 nm (4 voxel) distance
8 nm (5.3 voxel) distance
Overlay

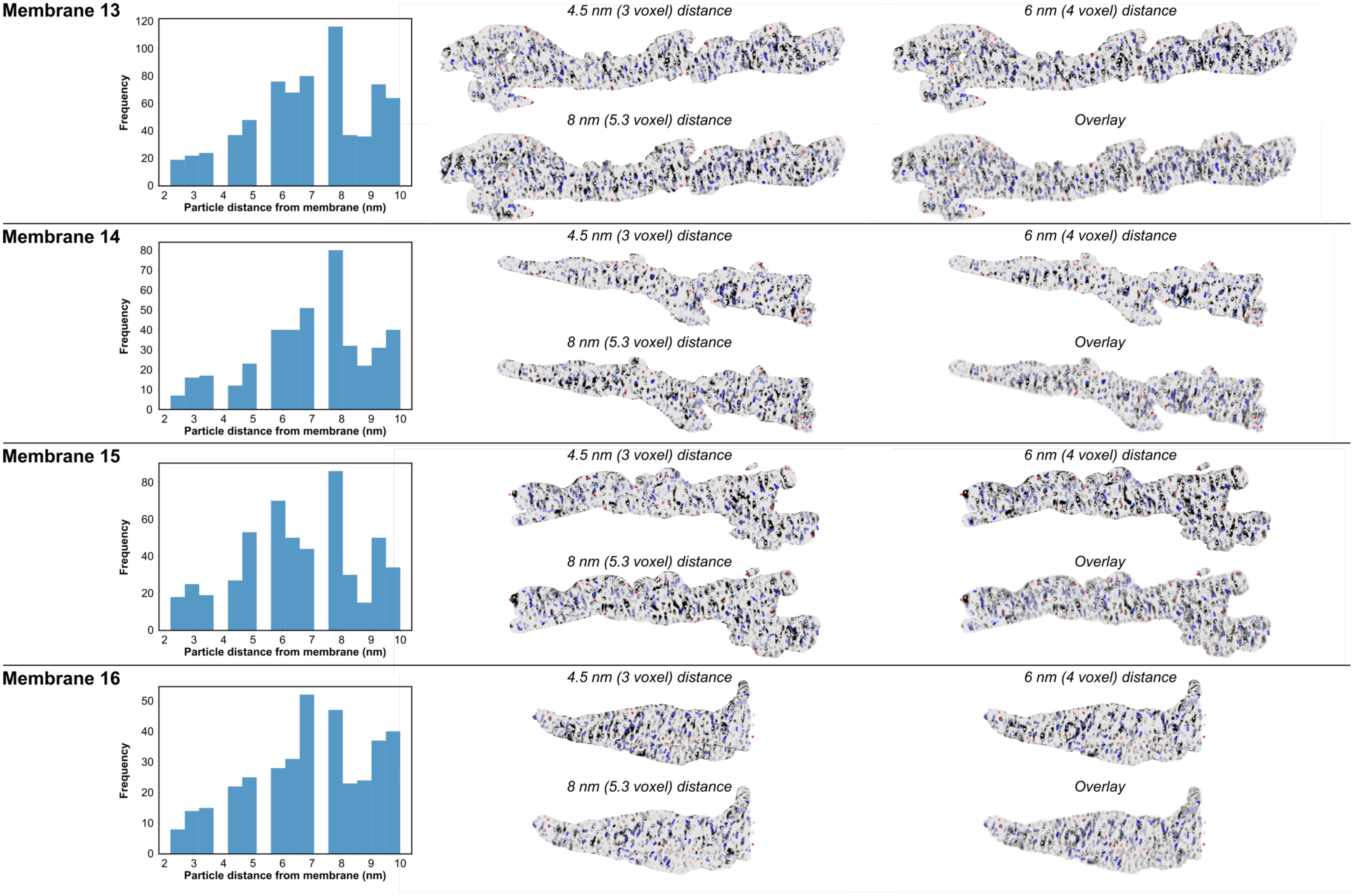
Membrane 13
Frequency
Particle distance from membrane (nm)
4.5 nm (3 voxel) distance
6 nm (4 voxel) distance
8 nm (5.3 voxel) distance
Overlay
Membrane 14
Frequency
Particle distance from membrane (nm)
4.5 nm (3 voxel) distance
6 nm (4 voxel) distance
8 nm (5.3 voxel) distance
Overlay
Membrane 15
Frequency
Particle distance from membrane (nm)
4.5 nm (3 voxel) distance
6 nm (4 voxel) distance
8 nm (5.3 voxel) distance
Overlay
Membrane 16
Frequency
Particle distance from membrane (nm)
4.5 nm (3 voxel) distance
6 nm (4 voxel) distance
8 nm (5.3 voxel) distance
Overlay

**Supplemental Figure 6** Membrane-linked protein coordinates identified by 'agnostic' template matching. Left - histograms represent distances of picks from segmented membrane surface. Right – 'en-face' surface density projections showing concordance of template matching picks with membrane densities. Projections are contoured to 4.5 nm, 6 nm and 8 nm from the segmented membrane surface.

**Supplemental Table 1** Summary of selected tomogram positions and their main features

| Tomogram | Features visible |
|---|---|
| **Position002** | ER, ribosomes |
| **Position003** | ER, ribosomes, vesicles |
| **Position004** | ER, ribosomes, vesicles, nuclear membrane, nuclear pore, chromatin |
| **Position005** | mitochondria, ER, ribosomes |
| **Position006** | microtubule organising center |
| **Position010** | mitochondria, ribosomes |
| **Position015** | mitochondria, ER, ribosomes |
| **Position016** | mitochondria, ribosomes |
| **Position022** | mitochondria, ribosomes |
| **Position023** | mitochondria, ER, ribosomes |
| **Position025** | ER, ribosomes |
| **Position029** | mitochondria, ER |
| **Position030** | ER, ribosomes, vesicles |
| **Position031** | ER, vesicles |
| **Position033** | mitochondria, ER, ribosomes |
| **Position035** | mitochondria, ribosomes |
| **Position036** | mitochondria, ribosomes, vesicles |
| **Position038** | ER, vesicles |
| **Position043** | mitochondria, ER, vesicles |
| **Position047** | ER, ribosomes, vesicles |
| **Position048** | ER, ribosomes |
| **Position050** | ER, vesicles |
| **Position051** | nuclear membrane, nuclear pore, ER, ribosomes |
| **Position054** | ER, ribosomes |
| **Position060** | ER |
| **Position062** | ER, Golgi, vesicles |
| **Position065** | mitochondria, ER, ribosomes |
| **Position073** | ER, ribosomes |
| **Position074** | ER, vesicles |
| **Position075** | nuclear membrane, ER, Golgi, vesicles |
| **Position076** | ER, Golgi, vesicles |
| **Position077** | ER, ribosomes, vesicles |
| **Position089** | nuclear membrane, nuclear pore, ER, ribosomes |
| **Position090** | ER, ribosomes, vesicles |
| **Position091** | ER, ribosomes |
| **Position095** | ER, ribosomes |
| **Position106** | mitochondria, ribosomes |
| **Position108** | mitochondria, ribosomes |
| **Position110** | ER, Golgi (x2) |
| **Position111** | mitochondria, ER, ribosomes |

| | |
|---|---|
| **Position112** | ER, ribosomes |
| **Position113** | Golgi, vesicles |
| **Position123** | mitochondria, ER, ribosomes |
| **Position124** | mitochondria, ER, ribosomes |
| **Position125** | mitochondria, ER, ribosomes |
| **Position127** | ER, vesicles |
| **Position147** | mitochondria, Golgi, ribosomes |
| **Position190** | ER, Golgi, ribosomes |
| **Position269** | ER |
| **Position271** | Golgi |
| **Position274** | Golgi |
| **Position278** | ER, ribosomes |
| **Position321** | ER, Golgi, TGN |
| **Position322** | ER, ribosomes, Golgi |
| **Position323** | ER |
| **Position326** | ER, ribosomes |
| **Position354** | ER, Golgi, ribosomes |

**Supplemental Table 2** Comparison of PyTME performance across templates used in this work

| Template | Picks across Position321 |
|---|---|
| **2AM3** | 260,015 |
| **5ZIB** | 185,851 |
| **7Q4I** | 201,915 |
| **7ZAY** | 79,713 |
| **8CCY** | 183,915 |
| **Spheroid** | 254,810 |
| **Spheroid after korpuskulum subtraction** | 13,740 |
